# Opposing transcriptomic gradients explain orthogonal maps in human visual areas

**DOI:** 10.1101/812560

**Authors:** Zonglei Zhen, Jesse Gomez, Kevin S. Weiner

## Abstract

Opposing transcriptomic gradients explain the large-scale organization of cortex. Here, we show that opposing transcriptomic gradients also explain the fine-scale organization of orthogonal maps in human visual areas. We propose a model relating transcriptomics, cell density, and function, which predicts that specific cortical locations within these visual maps are microanatomically distinct and differentially susceptible to genetic mutations. We conclude with histological and translational data that support both predictions.

## Maint Text

Active transcription of a small set of genes contributes to the large-scale arealization of functional hierarchies and gradients in human cortex. Two recent studies reveal that both the broad layout of functional regions^1^ and even the hierarchical ordering of regions within a processing stream^2^ are accurately described by genetic transcription that produces opposed, linear gradients in cortex. However, whether this broad relationship extends to smaller spatial scales is unknown. Thus, the goal of the present study was to answer the following fundamental question: *What role do genes play in the organization of functional gradients, or topographic maps, within a single human cortical area?*

While genetic gradients (e.g., Ephrins) during gestation establish the topography of connectivity in visual cortex^3^, the role that additional genes play in the layout of topographic maps within cortical areas in the adult brain is less well understood. To fill this gap in knowledge, we employed a dataset^4^ detailing the transcriptomic landscape of adult human cortex from the Allen Human Brain Atlas (AHBA). Early visual areas (V1, V2, V3) were used as target areas because they contain two, well-characterized orthogonal maps of eccentricity and polar angle^5^ (Fig 1a) that can be modeled in the absence of functional data^6^. Using tools from our previous work^2^, tissue samples and functional areas were aligned to the same cortical surface.

**Figure 1:**
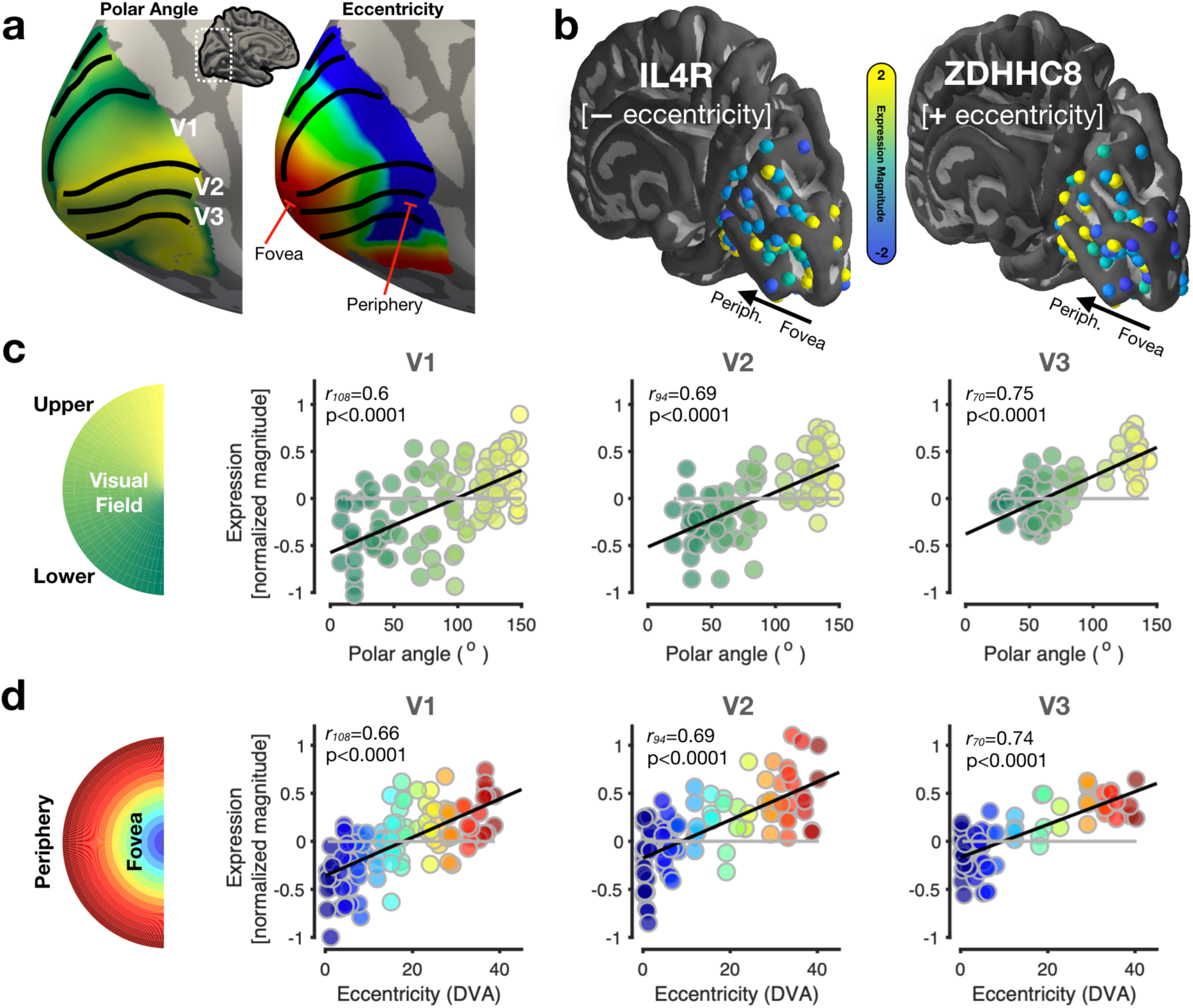
Opposing transcriptomic gradients explain orthogonal maps in human visual areas. **(a)** Example orthogonal maps of polar angle and eccentricity on an inflated cortical surface. Areas V1, V2, and V3 are defined (Online Methods). **(b)** Right hemisphere tissue samples of the AHBA projected onto the cortical surface. Inset bar displays colormap of normalized expression magnitude. Example genes whose transcription levels form either a negative gradient (left) or positive gradient (right) in relation to eccentricity. **(c)** Correlations between a tissue sample’s polar angle preference and the mean expression of the top 1% of genes. For illustration purposes, the positive and negative gradients have been combined by inverting the sign of the negative gradient. As illustrated in Fig S1, the same result occurs when calculating correlations separately for each gradient. Black line: Line of best fit. Gray line: Average correlation from a bootstrapping (N=1000) procedure in which genes were randomly selected (Online Methods). **(d)** Same as (c) but for the underlying eccentricity value of each tissue sample. DVA: degrees of visual angle.

We tested the *a priori* hypothesis that opposed linear gradients are not only a feature employed by cortex at the large-scale^1, 2^, but also at finer spatial scales within cortical areas. We identified tissue samples (n=136) from the AHBA located within human early visual areas that form linear gradients correlated with eccentricity or polar angle. To minimize false alarms, we selected only the top 1% of positively and negatively correlated genes ranked by the negative log of their p-values as in previous work^2^. Example genes whose expression correlated positively (higher expression in periphery) or negatively (lower expression in periphery) with increasing eccentricity are illustrated in Fig 1b. In each area, we observe significant correlations (r’s > 0.6, p’s < 0.001) between the magnitude of genetic transcription and either polar angle or eccentricity, suggesting that the orthogonal maps of receptive field (RF) properties are described by equally orthogonal genetic gradients, which is consistent with our hypothesis. All correlations were between 29.5 and 40.2 standard deviations above their respective bootstrapped (n=1000) null distributions (gray line in Fig 1c-d; Online Methods).

Interestingly, the identified genes are largely area- and map-specific. When calculating the dice coefficient between the genes contributing to the layout of eccentricity and polar angle within each area, there is little (∼3%) overlap (dice of V1: 0.12; V2: 0.07; V3: 0.07; see Fig S2 for overlap between areas). The identified genes (Table S1) are independent of those identified previously that contribute to the large-scale arealization of the visual processing hierarchy (Fig S2), indicating that cortical organization employs linear transcription gradients at multiple spatial scales using unique sets of genes. Additionally, polar angle and eccentricity measurements of one area (e.g., V1) do not predict (-0.33 < r’s < 0.34, p’s > 0.05; Figs S3-S6) the gene expression in another area (e.g., V2 or V3).

The latter finding is particularly surprising considering that the representation of eccentricity is shared across areas V1-V3. We hypothesized that the identified genes may have different roles in the development of features that contribute to functional differences between areas and within a given topographic map. Two parsimonious features fit these criteria. Firstly, RFs are smaller and denser in foveal compared to peripheral cortex, as well as smaller in V1 compared to V2 and V3 (Fig 2a)^7^. Secondly, negative eccentricity gradient genes code for many microstructural proteins (Fig 2b) whose expression also decreases from fovea to periphery. Together, denser RFs and microstructure-promoting transcription generate a functional-genetic model (Fig 2c) that makes the explicit prediction that human foveal cortex has an increased density of cells and relevant neuropil to support the increased fidelity of vision associated with foveal processing. Employing a new histological quantification approach (Fig 2d), we successfully demonstrate differences in cell/neuropil density between foveal and peripheral V1 (Fig 2d, 2e) for the first time in humans, which is consistent with measurements from non-human primates^8^.

**Figure 2:**
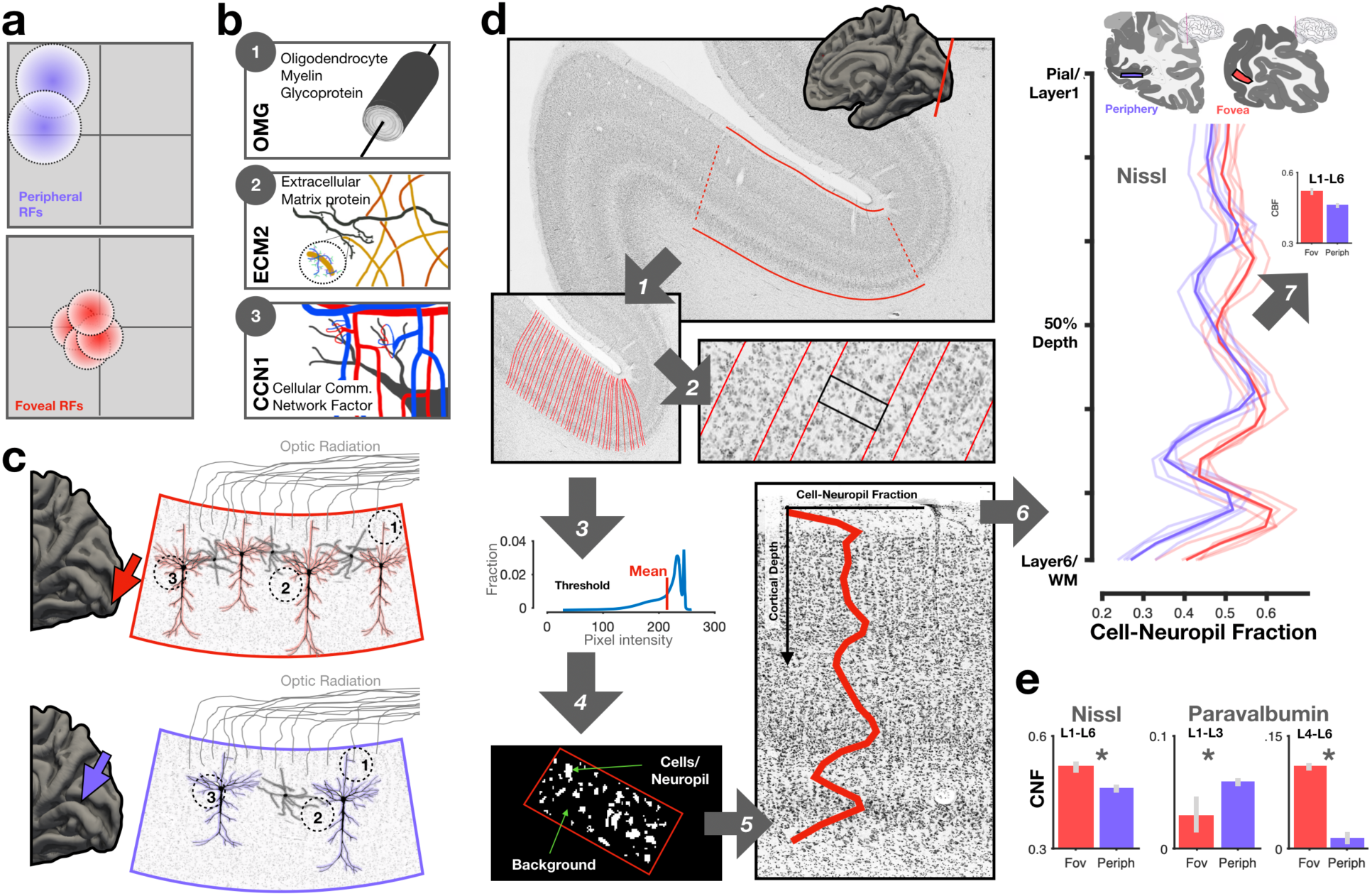
Validating tissue differences between foveal and peripheral cortex predicted by a functional-genetic model. **(a)** Schematic showing that receptive fields (RFs) are denser and smaller in foveal (red) compared to peripheral (purple) cortex. **(b)** Identified genes with increased expression in foveal compared to peripheral cortex (Table S1) suggests that higher RF density in foveal cortex is associated with higher expression of genes that regulate macromolecular tissue structures such as myelin (OMG), extracellular matrix material (ECM2; collagen, yellow curves; glycoproteins (blue/purple inset), and capillary junctions (CCN1). **(c)** A functional-genetic model resulting from (a) and (b) hypothesizes that higher RF density and increased expression of genes controlling key tissue structures will result in higher cellular/neuropil density in foveal cortex (posterior calcarine sulcus, red arrow) compared to peripheral (anterior calcarine sulcus, purple arrow). Right: Schematic illustration of cellular/ neuropil density differences between foveal (top) and peripheral (bottom) cortex. Colors: neurons. Gray: interneurons. Dotted circles (1-3): Where proteins of the genes in (b) are likely localized. **(d)** Basic steps of the approach developed here to quantify cytoarchitecture from coronal Nissl stains, which derives the fraction of pixels above background which we deem cell-neuropil fraction (CNF) along cortical traversals modeled to follow the organization of cortical columns. See Online Methods for description of the pipeline resulting in average CNF measurements across cortical layers (upper right inset) and histological slices. Red: foveal; Purple: peripheral slices of pericalcarine cortex. Dark contours represent the mean (step 6). Upper insets indicate the location of the histological slices from which each ROI (colored rectangles) was defined. **(e)** Left: Bar graph indicates that foveal portions have a significantly higher (p<.005) CNF (0.52 +/- 0.02) compared to peripheral portions (0.45 +/- 0.02) from Nissl stained sections. Right: Control analyses of paravalbumin stains show that CNF varied by cortical layer (Layers 1-3, peripheral > foveal, t(8)=2.72, p<0.03; Layers 4-6, foveal > peripheral, t(8)=5.2, p<0.001) indicating that foveal and peripheral cortical slices do not always vary by a global mean difference simply as a result of potential processing biases of the histological tissue (Online Methods).

This functional-genetic model also makes the translational prediction that mutations in a subset of the identified genes should produce specific visual field deficits, likely unevenly across a given map. We verify the feasibility of this prediction with CXorf58 (Fig 3), an X chromosome gene whose expression negatively correlated with eccentricity and has been linked to retinitis pigmentosa (RP), which results in perceptual deficits of the peripheral visual field. Mutations in CXorf58 may also explain recent findings^9^ showing that females with Turner Syndrome (TS) — a condition in which an X chromosome is damaged or missing^10^ — present with deficits in RF coverage of the peripheral visual field specifically in V2 and V3, where CXorf58 is negatively correlated with eccentricity (Fig 3). Additional mutations in SLITRK4 and RAI2 — both X chromosome genes involved in axonogenesis^11^ and cell growth^12^ and identified here — could adversely impact peripheral representations where tissue microstructure reductions would hinder the fidelity of neural circuits underlying the pooling of visual information.

**Figure 3:**
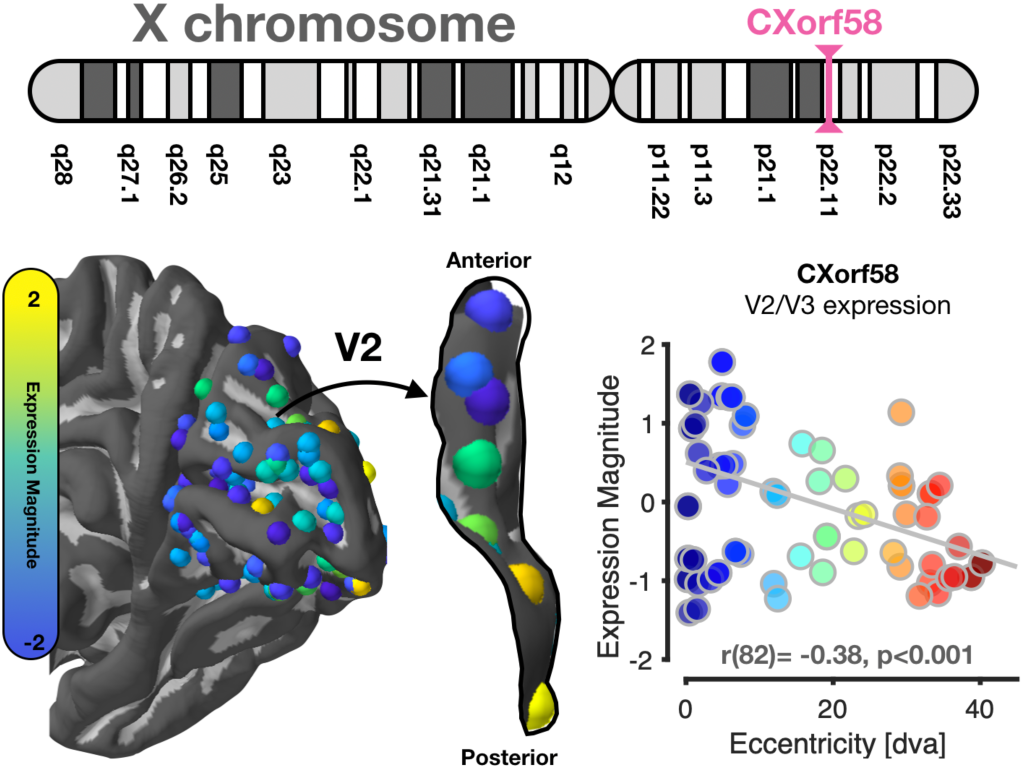
Translational insights into the role of genetic mutations in the loss of peripheral vision. The functional-genetic model makes the translational prediction that mutations in a subset of the identified genes should produce specific visual field deficits. *Top*: The location of CXorf58 on the short arm of the X chromosome. Mutations in CXorf58 have been linked to retinitis pigmentosa (RP), which results in peripheral vision deficits. *Bottom left:* Expression pattern of CXorf58 within the human occipital lobe. Expression within dorsal V2 is emphasized given recent findings showing that patients with Turner syndrome have reduced peripheral visual field coverage in V2 and V3 compared to controls. *Bottom right:* Expression pattern of CXorf58 negatively correlates with eccentricity. We propose that CXorf58 mutations may not only be linked to deficits associated with RP, but also Turner syndrome.

Altogether, our results suggest a new rule of cortical organization in which the adult brain employs opposed transcriptional gradients at multiple spatial scales: across the cortex^1^, across areas within processing hierarchies^2^, and now across maps within a single area. Previously, we hypothesized that large-scale transcription gradients contributed to the differences in population RFs across areas of the visual processing hierarchy. Here, we improve the resolution of this hypothesis by an order of magnitude to the anatomical scale of cellular structures and the functional scale of RFs within a single visual map. While transcriptomic gradients contribute to the layout of RF properties within and across field maps, additional evidence suggests that such transcription shows protracted development^2^, and that RFs are even sculpted through experience and viewing behaviors^13^. Thus, experience likely modulates the transcription of genes involved in structural mechanisms underlying RFs. Future work will add additional features such as connectivity^3^ and cortical layering^17^ to the present model, which will continue to deepen our understanding of the balance between transcriptomic and experiential contributions to the structure and function of human cortex.

## Online Methods

### Data

All analyzed data were curated from the following open datasets:

1. Human Brain Atlas (AHBA) Microarray Data: http://human.brain-map.org
2. atlas of human visual cortex: https://github.com/noahbenson/neuropythy
3. histological stains: http://www.brainspan.org/static/atlas

### Defining eccentricity and polar angle maps in areas V1, V2, and V3

Human visual areas are defined by two orthogonal functional maps: one of a receptive field’s (RF) polar angle describing its polar position along the circular image of the retina, and one of eccentricity, describing its distance from the center of the visual field (the fovea). While maps of eccentricity are shared across early visual areas, the transition of one area to another is based on differences in polar angle^5^. Polar angle transitions between early visual areas occur with such regularity relative to the folding of cortex that a model based on an individual’s cortical anatomy is sufficient to automatically define areas V1 through V3 as precisely as manual definitions of these areas^6^. We applied this model to a cortical surface of the MNI152 brain, which is an anatomical average of human cortex from the Montreal Neurological Institute made from the average of 152 brains. This cortical space was chosen given that tissue samples from the donor brains of the AHBA have been aligned to the same MNI volumetric space. A cortical surface was produced for this anatomical volume using FreeSurfer^14^.

### AHBA Tissue Samples

The AHBA provides transcription data across human cortex, pooled from six donors and processed via Microarray. Microarray samples underwent strict normalization and data quality checks through AHBA as described here: http://human.brain-map.org. For data organization, preprocessing was identical to our previous work^2^. Importantly, we follow the standardized processing pipeline proposed by Arnatkeviciute, Fulcher, and Fornito^15^. Briefly, the raw microarray expression data for each of the six donor brains included the expression level of 29,131 genes profiled via 58,692 microarray probes. We implemented five preprocessing steps. First, probes were excluded that did not have either a) a gene symbol or b) an Entrez ID. This resulted in 20,737 genes. Second, the expression profiles of all the probes targeting the same gene were averaged. Third, in order to remove variability in gene expression across donors, gene expression values of tissue samples were normalized by calculating z-scores separately for each donor. Fourth, because previous studies did not identify significant inter-hemispheric transcriptional differences, data from both hemispheres were combined. Finally, comparing a tissue sample’s genetic expression to either the eccentricity or polar angle of its underlying cortex required mapping these tissue samples to a common cortical space. Each sample from the AHBA was associated with a 3D coordinate from its donor’s MRI brain volume and its corresponding coordinate (*x*, *y*, *z*) in the MNI152 space. Each tissue sample was mapped to the nearest cortical surface vertex with a 5mm distance threshold and assigned to the visual field map in which that surface vertex was contained. This resulted in the following number of tissue samples aligned to each visual area: 54 in V1, 47 in V2, and 35 in V3.

### Comparing gene transcription and visual field map properties

With tissue samples and maps of eccentricity and polar angle having been mapped to the same cortical surface, we extracted the underlying eccentricity or polar angle from each tissue sample by taking the average of the vertices that were within a one voxel radius of the vertex to which the tissue sample had been mapped. As our previous study identified that genes distinguishing regions along a cortical processing hierarchy are organized into opposed, linear gradients^2^, we first tested the *a priori* hypothesis that opposed gradients would also exist within a single brain region to explain the orthogonal maps of eccentricity and polar angle of visual field maps. To that end, for each gene, we calculated the Pearson correlation between its expression magnitude across tissue samples within an area versus the mean eccentricity or polar angle of each tissue sample. Genes were then ranked by the negative log of their resulting p-values. As in the previous study^2^, the top 1% of positively correlated and the top 1% of negatively correlated genes were kept. The average expression value of those top genes are illustrated in Fig 1. This was done separately for each visual field map. For example, the mean expression of the top 1% (n=200) of positively correlated genes in V1 is averaged within each tissue sample and then plotted against its eccentricity or polar angle (pink circles of Fig S1). For purposes of visualization, we combined the two gradients together into a single gradient by flipping the sign of the negative gradient for Fig 1, but the opposed gradient with their original signs are shown in Fig S1. A list of the top positive and negative gradient genes for eccentricity and polar angle are listed in Table S1 for each visual field map. By averaging across the top genes, we aim to capture a general pattern that is shared across many genes, rather than focusing on a single gene which is likely more susceptible to a Type II error. As a control analysis, we sought to generate a null distribution for each correlation using a bootstrapping approach. Across n=1000 bootstraps, we repeated each correlation in each field map but randomly selected the same number of genes as the main correlation (from the total population of 20,737 genes). We plot the averaged correlation in Fig 1 as gray lines for each field map. We calculated, in units of standard deviation, the distance that the main correlations were from the bootstrapped null distribution. These distances were very large, ranging from 29.5 to 40.2 standard deviations, underscoring the significance of the observed correlations.

### Comparing inter-regional and intra-regional gene expression

In a previous study^2^, we identified n=200 genes that formed opposed positive (“ascending”) and negative (“descending”) gradients along the visual processing hierarchy of occipito-temporal cortex. We hypothesized that given that these genes were identified as those that distinguish regions within a processing stream, they should be unique, or non-overlapping, with the genes identified in the present study that are correlated with functional topography within a visual area. From these 200 inter-regional genes, and the intra-regional genes identified in each field map (2 orthogonal maps * 2 gradients * 200 genes per gradient = 800 genes per field map), we calculated the dice coefficient to describe the overlap of these two sets per field map. The dice coefficient (DC) is calculated as twice the number of overlapping genes normalized by the size of both lists. In this way, the highest DC possible for the dataset at hand is 0.5. The resulting DC for this analysis for V1, V2, and V3 is shown in Fig S2a. Of these 1000 total genes going into the DC analysis, approximately 2 overlapped. We also repeated this analysis for the list of intra-regional genes identified in two visual field maps. The resulting three DCs from the possible combination of comparisons for the three visual field maps is shown in Fig S2a. Next, to explicitly validate that the inter-regional genes identified in the previous study were not correlated with eccentricity or polar angle, separately for each of the “ascending” and “descending” gene gradients, we correlated the mean expression of these genes in the current tissue samples with eccentricity and polar angle in V1, V2, and V3. All of the resulting 12 correlations illustrated in Fig S2b were indeed non-significant: -0.2 < r’s < 0.22, p-values > 0.18. Lastly, we wanted to explicitly test if the genes identified from one region were correlated with functional properties in another region (Figs S3-S6). To do this, for example, we took the genes correlated with eccentricity in V2 and V3, removed the shared genes with V1 (e.g., genes that were also correlated with eccentricity in V1 were excluded) and then we correlated the expression of these unique V2/V3 genes in tissue samples of V1 with their underlying eccentricity. For all cases, we find no correlation (Bonferroni correction for multiple comparisons) with the exception of V2-unique genes being correlated with eccentricity in the tissue samples of V1.

### Quantifying histological slices of human cortex with BrainWalker

In order to test the feasibility of the hypothesis that gene transcription differences between foveal and peripheral representations within a single visual field map might induce microanatomical differences in the underlying cortical sheet, we developed a method to quantify human cytoarchitecture that a) was repeatable across different histological samples, b) could capture anatomical variation across cortical layers, and c) was importantly, observer-independent. To achieve this goal, we developed an analysis pipeline called BrainWalker that quantifies the fraction of cell bodies and neuropil (cell-neuropil fraction, CNF) above background noise in a given histological stain. Importantly, this software slides or “walks” across cortical layers and columns, producing a vectorized description of the CNF for a desired piece of cortical ribbon. The software works as follows, and is illustrated in Fig 2d. First, the user selects a ribbon of cortex to be quantified, tracing the middle of Layer 1 of cortex, and the corresponding length of the gray-white matter boundary (Fig 2d, box 1). The two boundaries are then split into 200 equally spaced points. Points in Layer 1 and the white matter boundary are modeled as positive and negative electric charges, respectively, and then equipotential lines are drawn connecting each corresponding point on the two boundaries (Fig 2d, arrow 1). These equipotential traversals are meant to model the shape of cortical columns traversing the cortical sheet, importantly fanning at gyral crowns^16^. Each traversal is then split into 30 equally spaced points, and then a window slides between two neighboring traversals along these 30 bins (Fig 2d, arrow 2). The underlying image (e.g., a Nissl stain) is thresholded at the mean pixel value to identify the “peaks” or puncta associated with cell bodies and neuropil structures (Fig 2d, arrow 3). While this threshold was somewhat arbitrary, a threshold was necessary to exclude irrelevant image structures such as holes from veins, tears in the tissue, or air bubbles on the mounting slide. These irrelevant values, or “background”, where the white slide mount becomes visible can be seen as the large peaks in the histograms of Fig S8. Fig S8 also shows that the chosen threshold does not impact the interpretation of the findings: histograms of all pixel values from the foveal and peripheral slices indicate that foveal slices (red) have greater tissue content at the entire range of lower pixel values corresponding to cortical tissue all the way up to the mean pixel intensity (solid black line) which marks the beginning of brighter pixels representing the slide background. Differences between foveal and peripheral slices at these higher pixel intensities result simply from the cropping of the images; foveal slices had more of the underlying slide visible as the posterior slices of the occipital pole were smaller than more anterior slices. After this thresholding, at every bin, the CNF (fraction of pixels belonging to cell bodies and neuropil versus background) is quantified (Fig 2d, arrow 4). The 30-bin vector for each traversal is then averaged across traversals, producing an average CNF contour detailing CNF from Layer 1 of cortex to white matter (Fig 2d, arrow 5). In Fig 2d, an example CNF contour is overlaid on a Nissl-stained section of human V1; one can appreciate how the peaks and valleys of the CNF contour highlight the different cell body densities across and between layers.

To generate CNF contours from human visual cortex, we used the freely available, high-resolution, Nissl-stained and Paravalbumin-stained coronal slices of human visual cortex from the Allen Institute’s BrainSpan Atlas (http://www.brainspan.org/static/atlas). In order to compare foveal and peripheral cortex, where genetic transcription is most differentiated in the present data and resulting anatomical differences are likely to be largest, we extracted slices (n=5) from the posterior Calcarine sulcus where foveal representations are located, and slices from the anterior Calcarine sulcus where the peripheral visual field is represented. To ensure that differences in overall stain intensity would not bias results, and to mirror the normalization steps for the transcriptional data, each slice was normalized by the mean pixel value of all the neighboring white matter pixels. On a region of the cortical ribbon spanning 1.25 centimeters, we extracted the CNF of each slice from V1 and plotted the CNF of each slice in Fig 2 comparing fovea and periphery for Nissl-stained cortex. We repeated this analysis with paravalbumin-stained cortex in Fig S6 (Fig 2e). For Nissl stains, we computed a simple two-sample t-test comparing the CNF between foveal V1 and peripheral V1 averaged across cortical layers (Fig 2e). For the paravalbumin stains, CNF varied by cortical layer, and t-tests were run comparing Layers 1-3 and Layers 4-6 separately (Fig 2e). Paravalbumin was analyzed not only as an additional feature that could vary by eccentricity, but as a control to ensure that foveal and peripheral cortical slices do not always vary by a global mean difference simply as a result of potential processing biases from the AHBA.

### Intra-regional genes on the X chromosome

Turner syndrome is a condition affecting females in which the individual is either missing an X-chromosome (monosomy X) or one of the X-chromosomes is damaged. While the condition is hallmarked by a number of physical atypicalities, genes on the short arm of the X chromosome (e.g., short stature homeobox gene, SHOX) have been linked to the stunted bone and limb growth that characterizes the small stature of those with Turner syndrome. Recent research has demonstrated that females with Turner syndrome present with visual deficits in the coverage of the periphery^9^. Retinotopic mapping of population receptive fields (pRF) in these individuals using functional MRI revealed that in V2 and V3, Turner females show reduced pRF coverage of the periphery compared to control females. With this finding in mind, we performed a directed search for genes in V2 and V3 that correlated with eccentricity located on the X chromosome, with special attention for those on the short arm of the chromosome. Table S2 shows the complete list of genes from V2 and V3 that were correlated with eccentricity and were located on the X chromosome. In Fig 3, we highlight gene CXorf58 (Chromosome X open reading frame 58), which is not only located on the X chromosome short arm, but is also a gene whose mutations are known to result in Retinitis Pigmentosa, which is characterized by the loss of peripheral vision. Other notable genes were SLITRK4, directly involved in regulating neurite outgrowth and arborization, as well as RAI2, involved in cell growth. Both of these genes likely contribute to the CNF differences we observe in typical adult cortex, and their mutation likely has especially adverse impact on peripheral representations where cell density is already lower compared to foveal cortex.

---

**Supplementary Figure 1:**
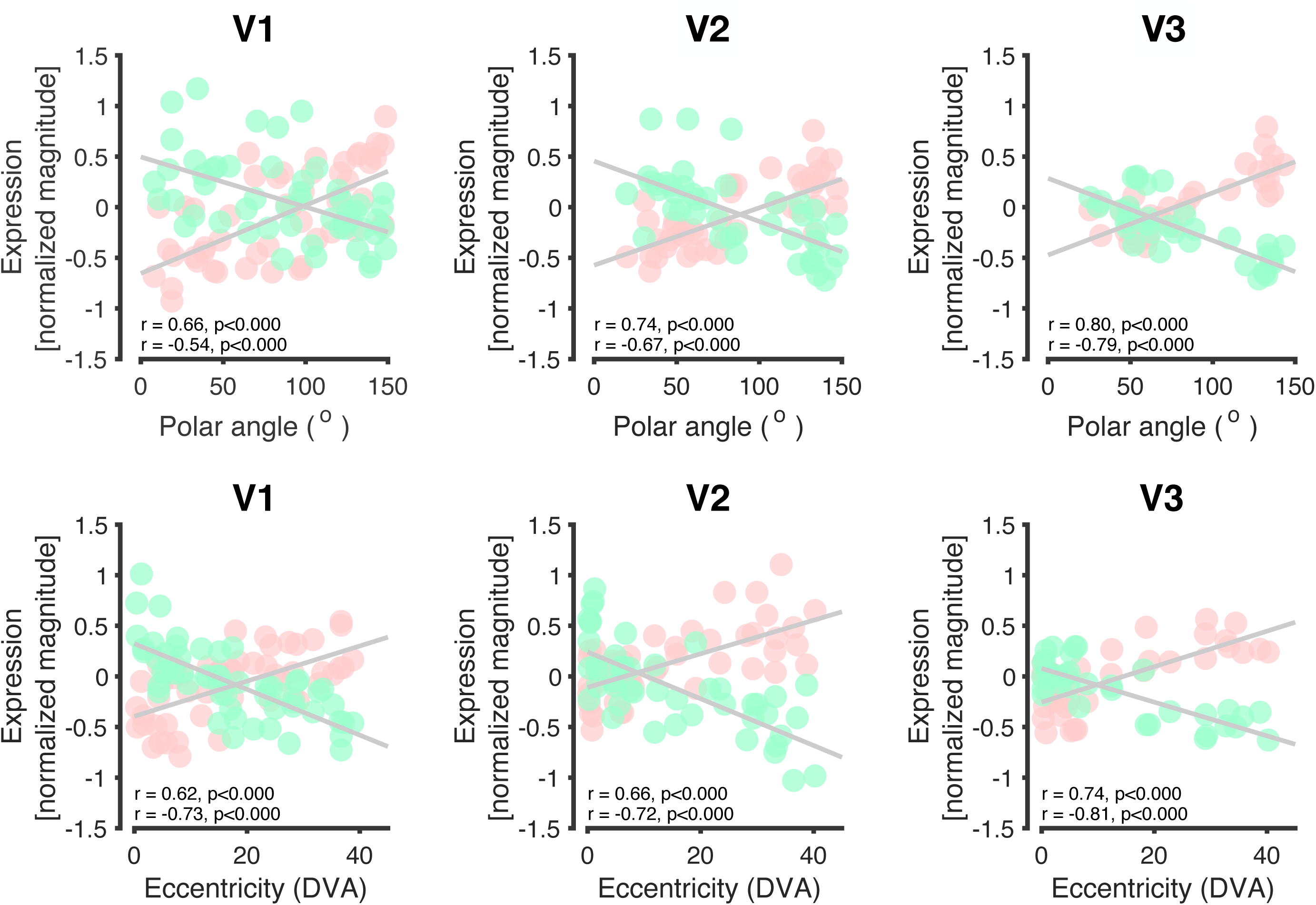
Correlations between functional representation (polar angle and eccentricity) and the expression of positive and negative gradient genes. The mean expression of positive gradient genes within tissue samples are represented in pink dots, while the mean expression magnitude of negative gradient genes are shown in green. Pearson correlations and the resulting p-values are shown as inset text in each plot. Line of best fit for the positive and negative correlations are overlaid as gray lines. For each gradient, there are the following number of tissue samples: V1 (n=54), V2 (n=47), V3 (n=35). (a) Polar Angle. (b) Eccentricity.

**Supplementary Figure 2:**
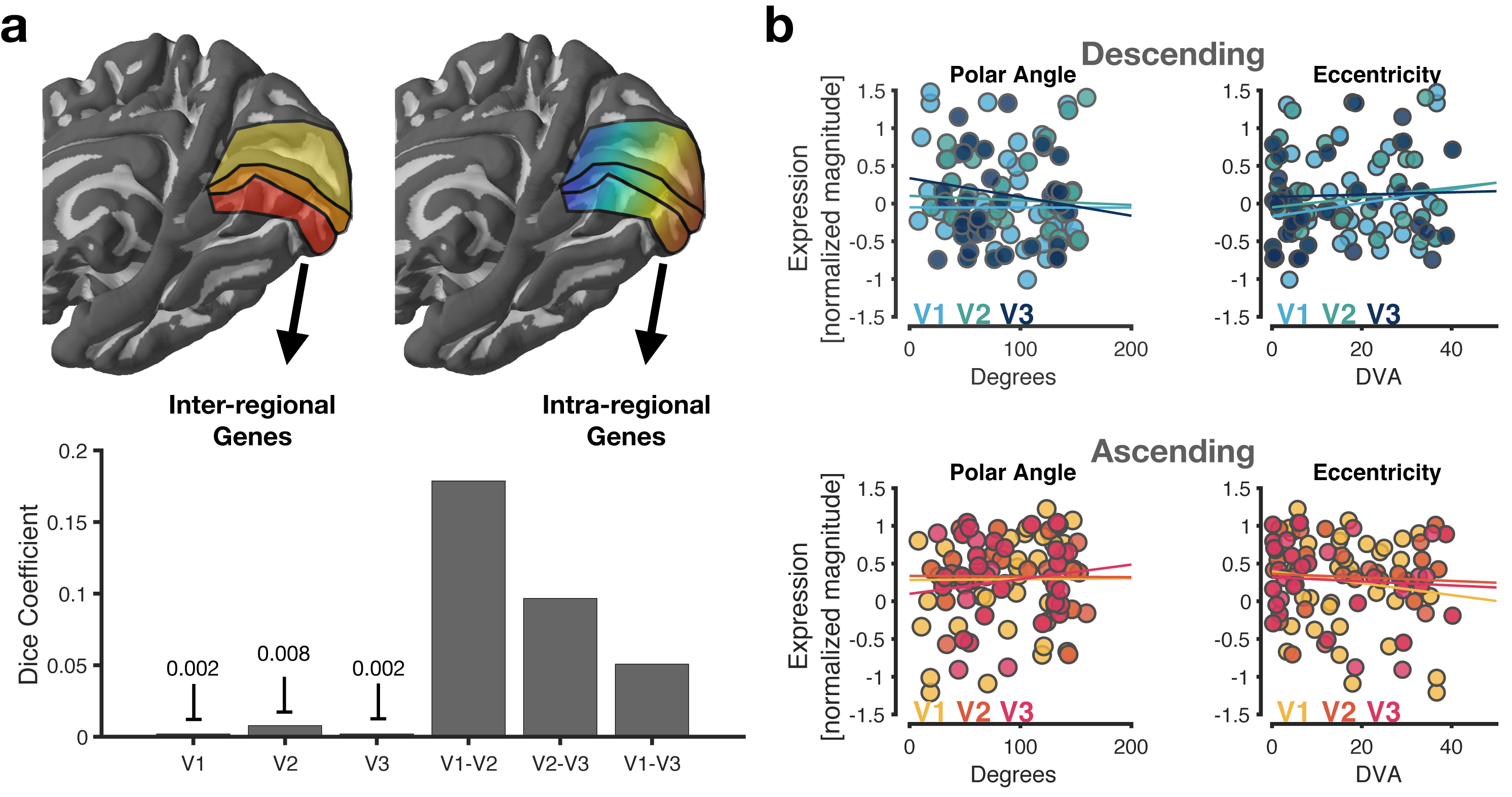
Genes involved in hierarchical ordering of the ventral visual processing stream (inter-areal genes) are distinct from those correlated with eccentricity or polar angle (intra-areal genes). (a) Left: Dice coefficients summarizing the overlap of intra-areal genes in V1 through V3 with previously identified inter-regional genes that distinguish regions across the visual processing hierarchy. Right: The overlap of intra-regional genes of one visual area relative to the others. V1 and V2 share the most genes. Overall, the intra-regional genes are largely restricted to a given area. (b) Mean gene expression levels relative to polar angle (left) or eccentricity (right) values for the previously identified inter-regional genes belonging to the descending (top) and ascending (bottom) gradients (-0.2 < r’s < 0.22, p-values > 0.18). That is, there were two clusters of genes identified in Gomez, Zhen, & Weiner 2019 that varied linearly in expression level as one traversed the visual processing hierarchy (e.g., from V1 to ventral temporal cortex): one cluster that ascended in expression level (positive correlation) and one that descended (negative correlation) in expression level from early to late positions of the visual processing hierarchy. The expression of these inter-regional genes for either the descending (top) or ascending (bottom) gradient were not correlated with polar angle or eccentricity within areas V1, V2, or V3.

**Supplementary Figure 3:**
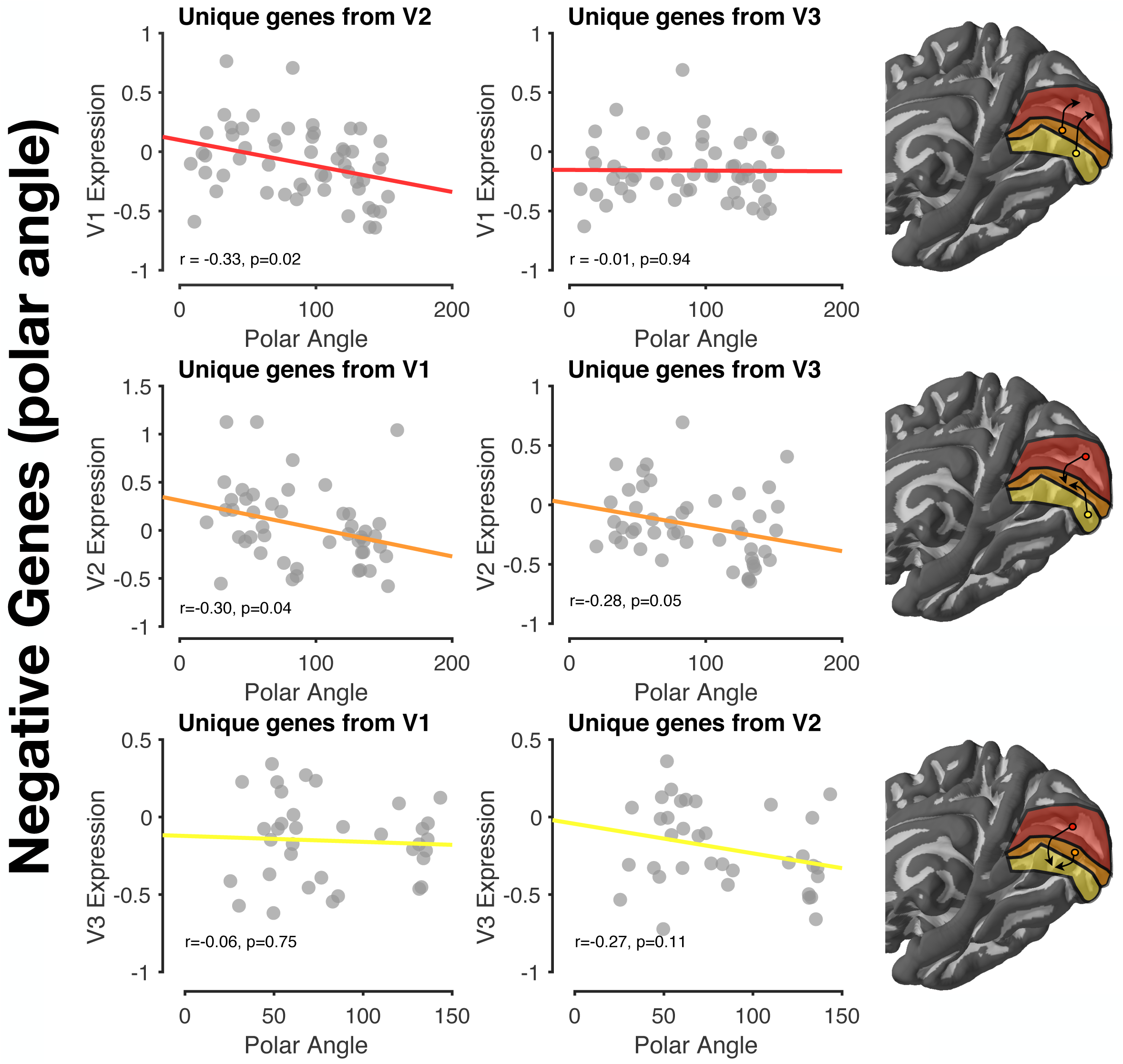
Correlations between polar angle and expression of negative genes from neighboring field maps. Polar-angle-correlated genes from each field map, while slightly overlapping, were largely a unique set of genes for each field map. To test whether the expression of genes from neighboring field maps were correlated with polar angle in the left out map, we took, for example, the genes from V2 and V3 correlated with polar angle, removed the overlapping genes that were correlated with polar angle in V1, and then correlated the mean expression of the remaining set of genes with polar angle in V1. The line of best fit is shown in each plot, and the resulting Pearson and p-values are inset as black text. Each gray circle is a tissue sample within a given field map.

**Supplementary Figure 4:**
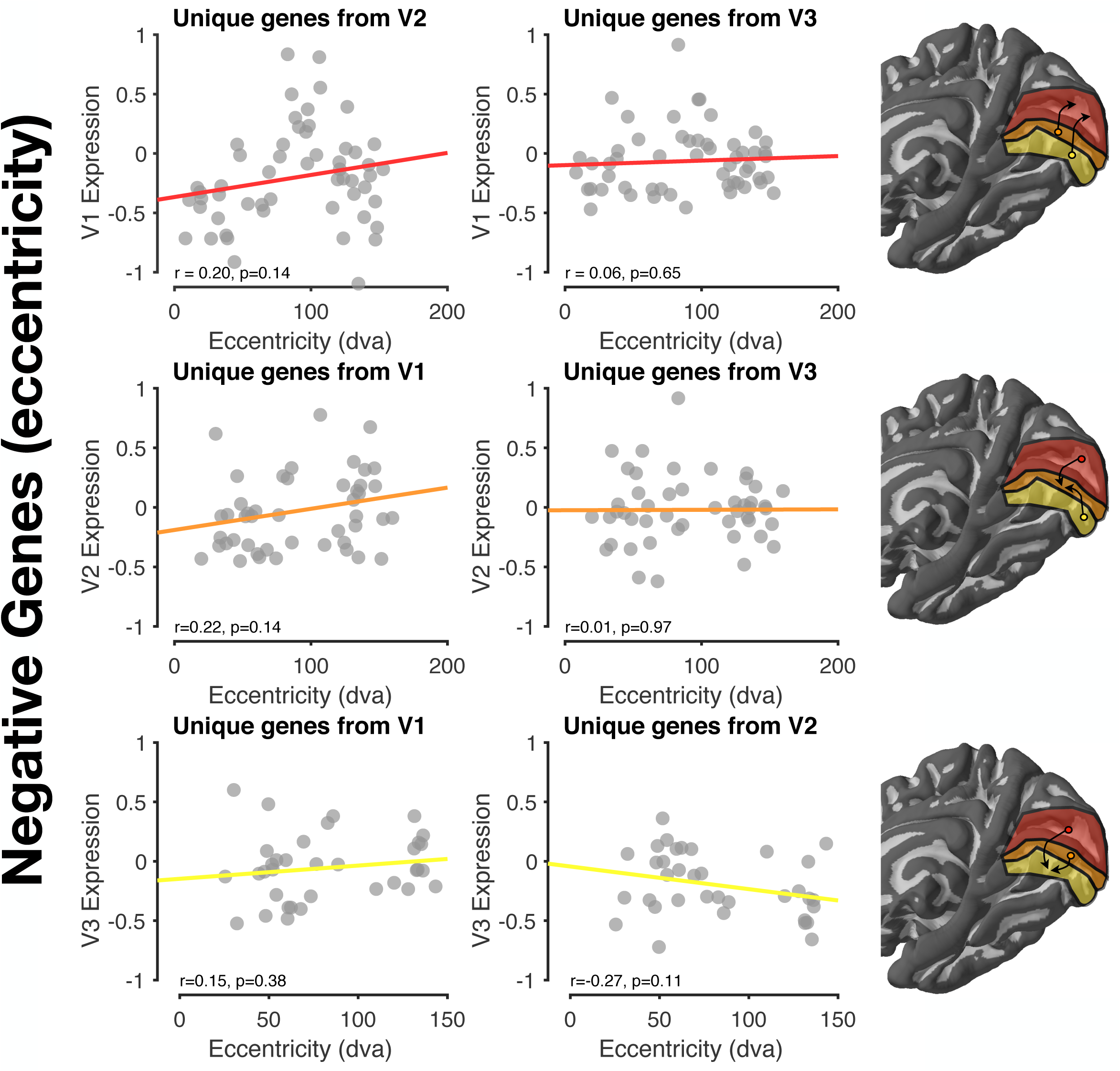
Correlations between eccentricity and expression of negative genes from neighboring field maps. Eccentricity-correlated genes from each field map, while slightly overlapping, were largely a unique set of genes for each field map. To test whether the expression of genes from neighboring field maps were correlated with eccentricity in the left out map, we took, for example, the genes from V2 and V3 correlated with eccentricity, removed the overlapping genes that were correlated with eccentricity in V1, and then correlated the mean expression of the remaining set of genes with eccentricity in V1. The line of best fit is shown in each plot, and the resulting Pearson and p-values are inset as black text. Each gray circle is a tissue sample within a given field map.

**Supplementary Figure 5:**
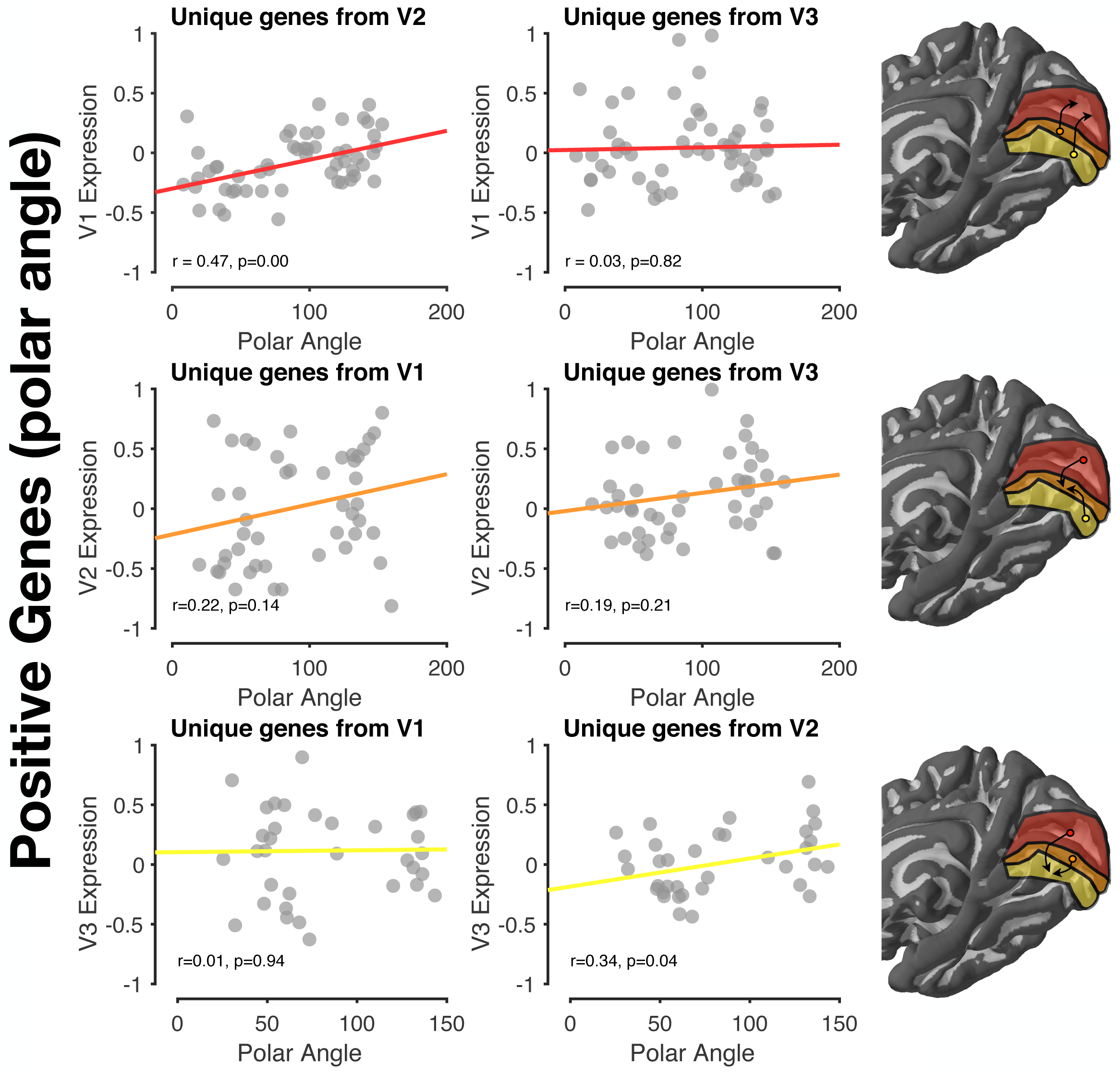
Correlations between polar angle and expression of positive genes from neighboring field maps. Polar-angle-correlated genes from each field map, while slightly overlapping, were largely a unique set of genes for each field map. To test whether the expression of genes from neighboring field maps were correlated with polar angle in the left out map, we took, for example, the genes from V2 and V3 that were correlated with polar angle, removed the overlapping genes that were correlated with polar angle in V1, and then correlated the mean expression of the remaining set of genes with polar angle in V1. The line of best fit is shown in each plot, and the resulting Pearson and p-values are inset as black text. Each gray circle is a tissue sample within a given field map.

**Supplementary Figure 6:**
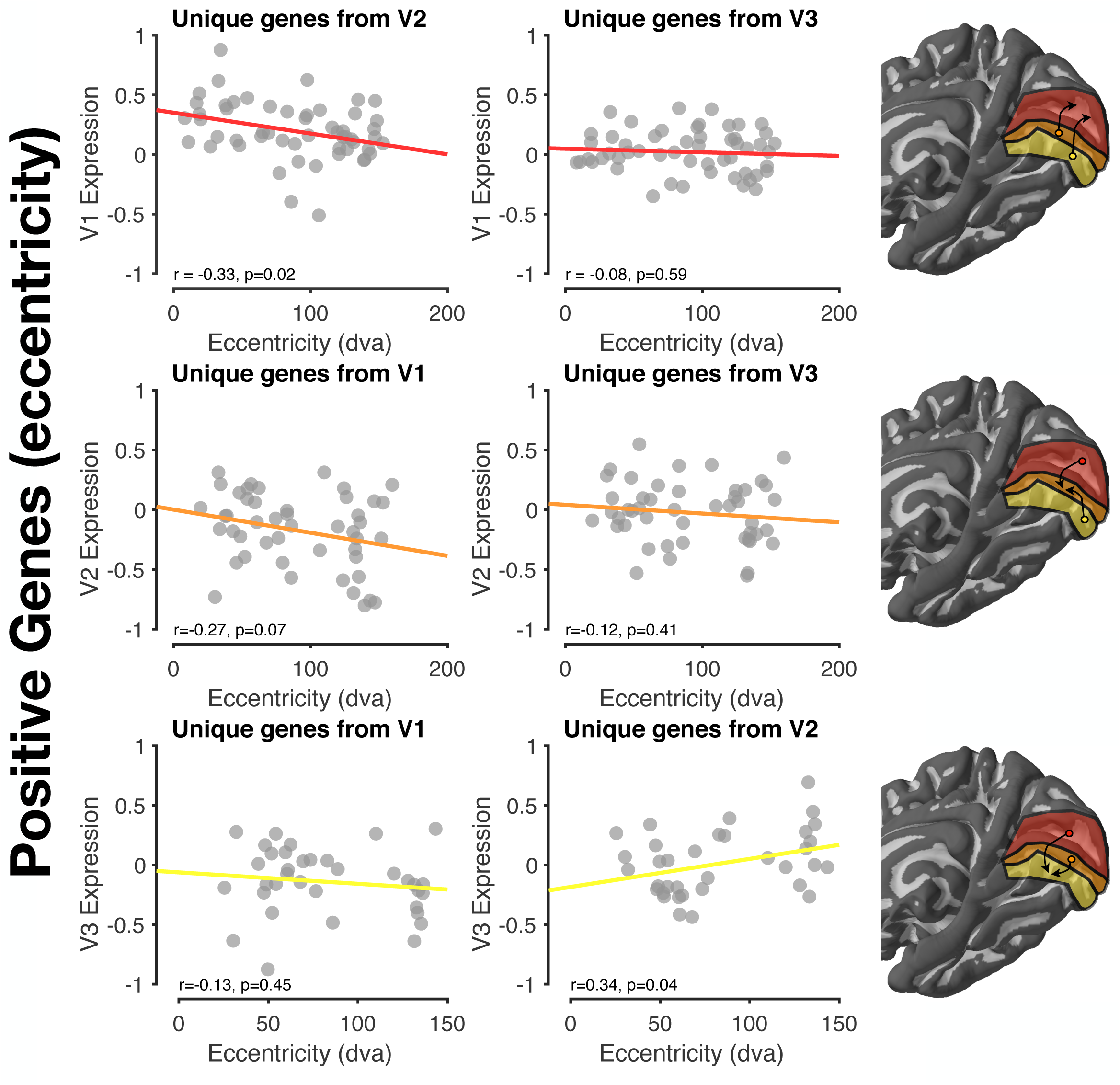
Correlations between eccentricity and expression of positive genes from neighboring field maps. Eccentricity-correlated genes from each field map, while slightly overlapping, were largely a unique set of genes for each field map. To test whether the expression of genes from neighboring field maps were correlated with eccentricity in the left out map, we took, for example, the genes from V2 and V3 that were correlated with eccentricity, removed the overlapping genes that were correlated with eccentricity in V1, and then correlated the mean expression of the remaining set of genes with eccentricity in V1. The line of best fit is shown in each plot, and the resulting Pearson and p-values are inset as black text. Each gray circle is a tissue sample within a given field map.

**Supplementary Figure 7:**
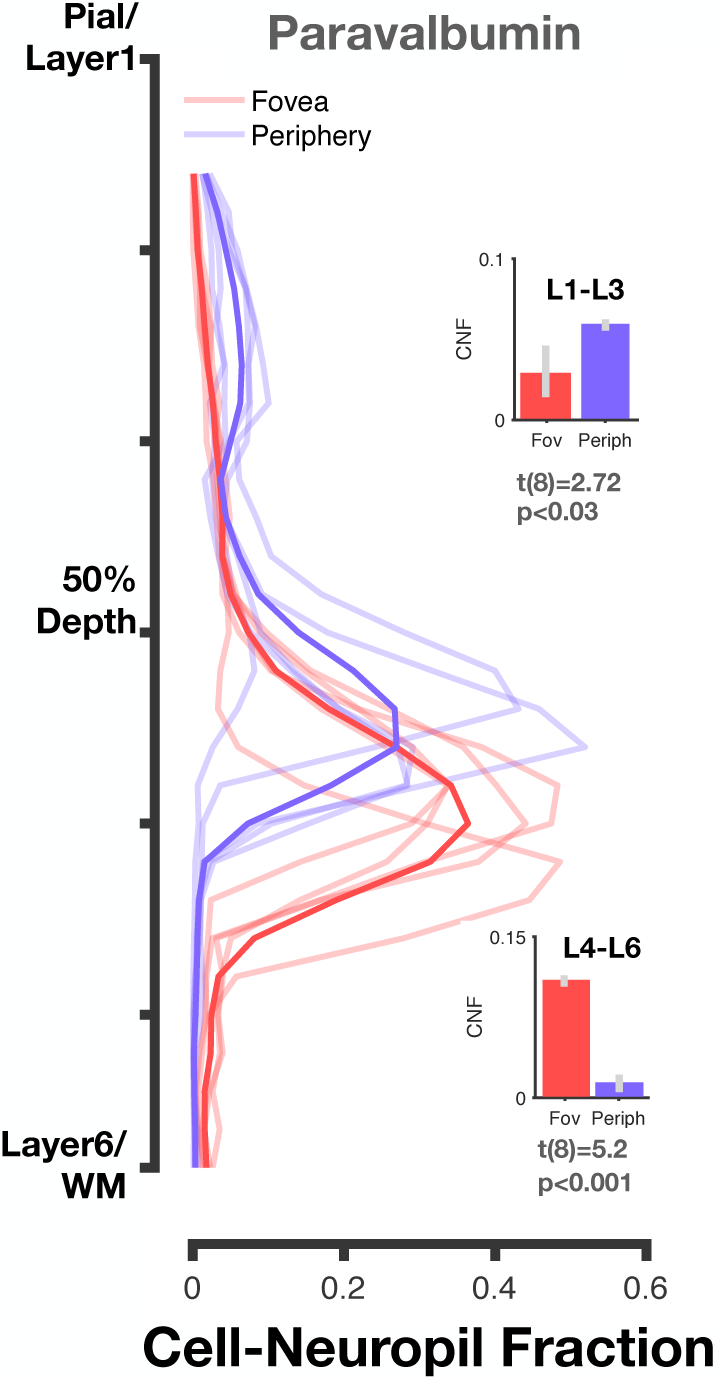
Differential presence of paravalbumin between foveal and peripheral visual cortex. CNF contours stained for Paravalbumin interneurons. We find that paravalbumin varies significantly between foveal and peripheral V1, with lower CNF in layers L4-L6 of peripheral cortex, and higher CNF in superficial layers (L1-L3). This analysis was conducted to verify that foveal and peripheral slices of V1 did not always vary by a global mean shift (potentially reflecting processing biases from AHBA).

**Supplementary Figure 8:**
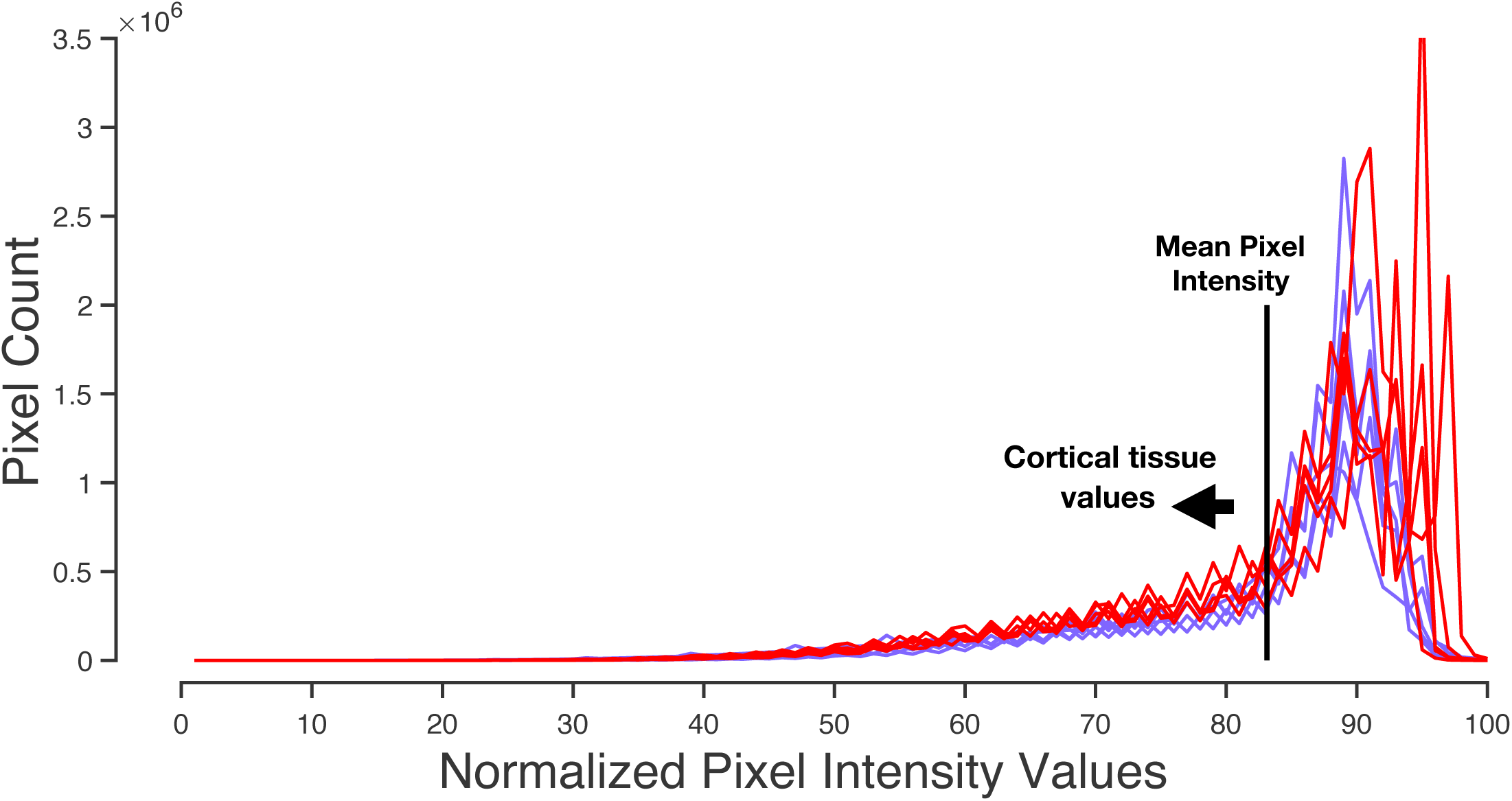
Histograms of pixel values from foveal and peripheral Nissl stains of human tissue. To ensure that our chosen threshold for the BrainWalker CNF analysis was not impacting our results, we plotted the histogram of all pixel values from the foveal (red) and peripheral (purple) Nissl-stained slices from the AHBA. Higher values on the x-axis represent brighter pixels, with the large peaks from around x=85 to x=100 corresponding to the white background of the images, which represent the slide on which the tissue was mounted. The mean pixel intensity (the threshold used for the CNF analysis) is shown with a solid black line. The reader can appreciate that at the entire meaningful range (>60) of the lower intensity values below the mean corresponding to cortical tissue, foveal slices (red) show higher values than peripheral (purple) slices.

**Supplementary Table 1:**
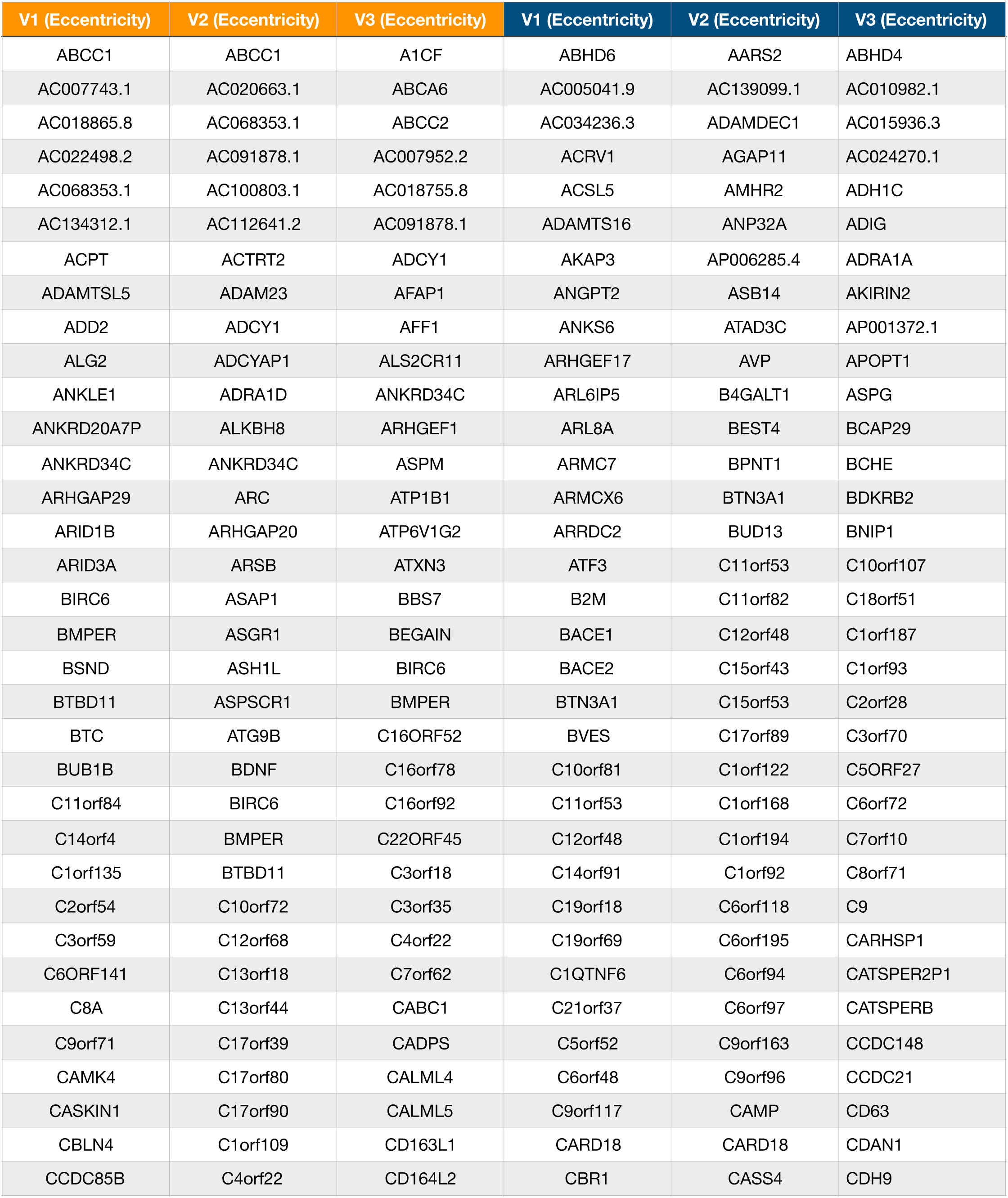
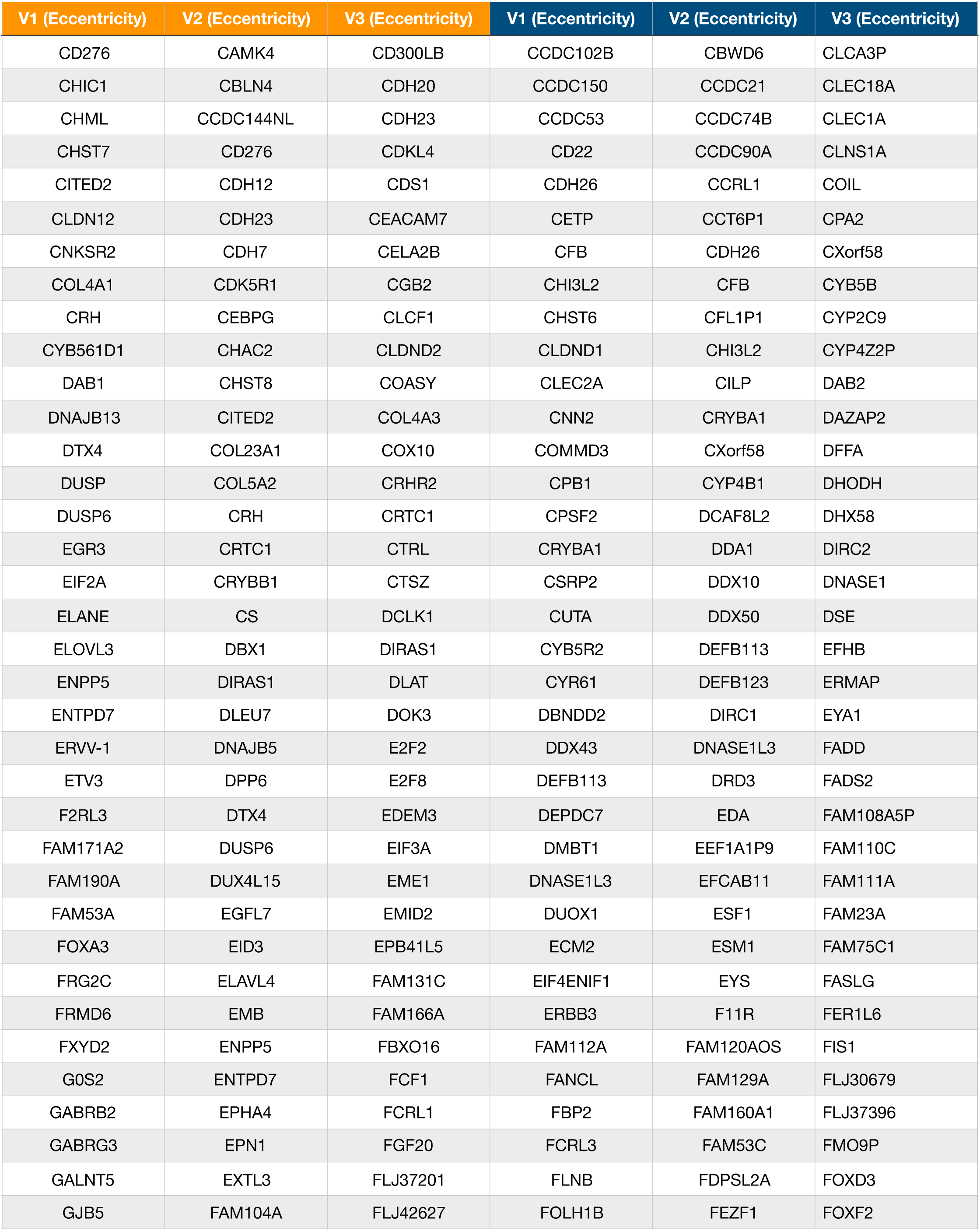
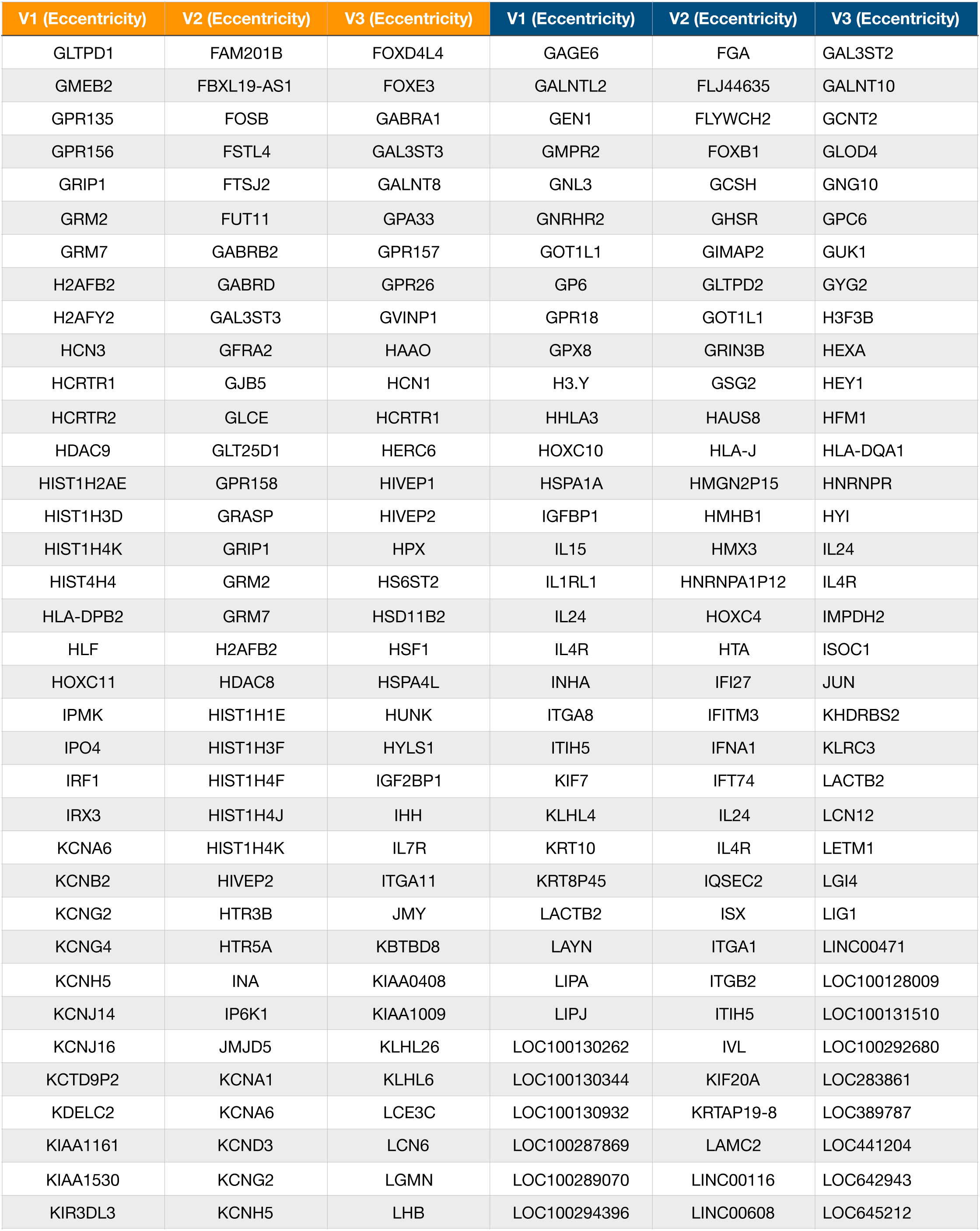
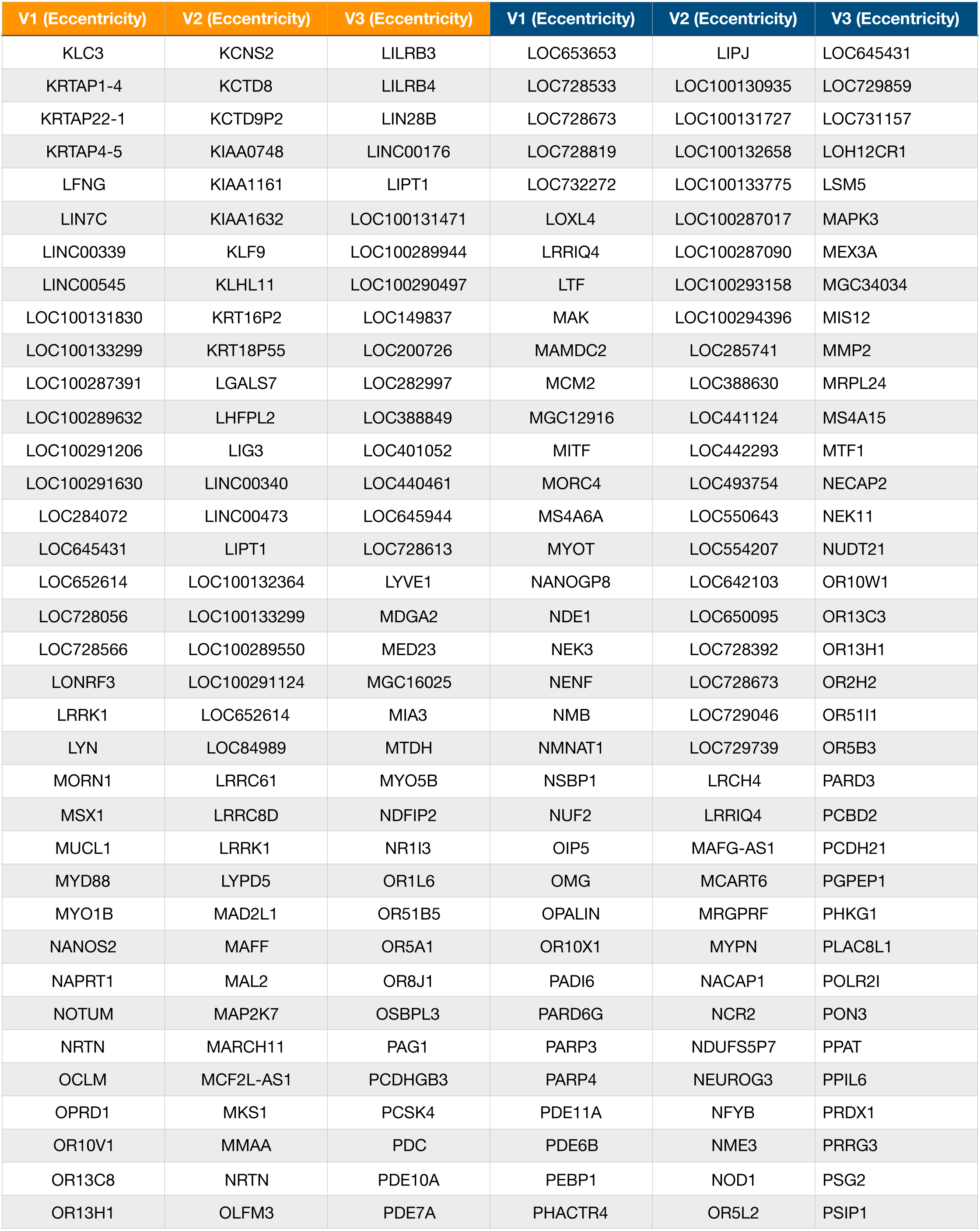
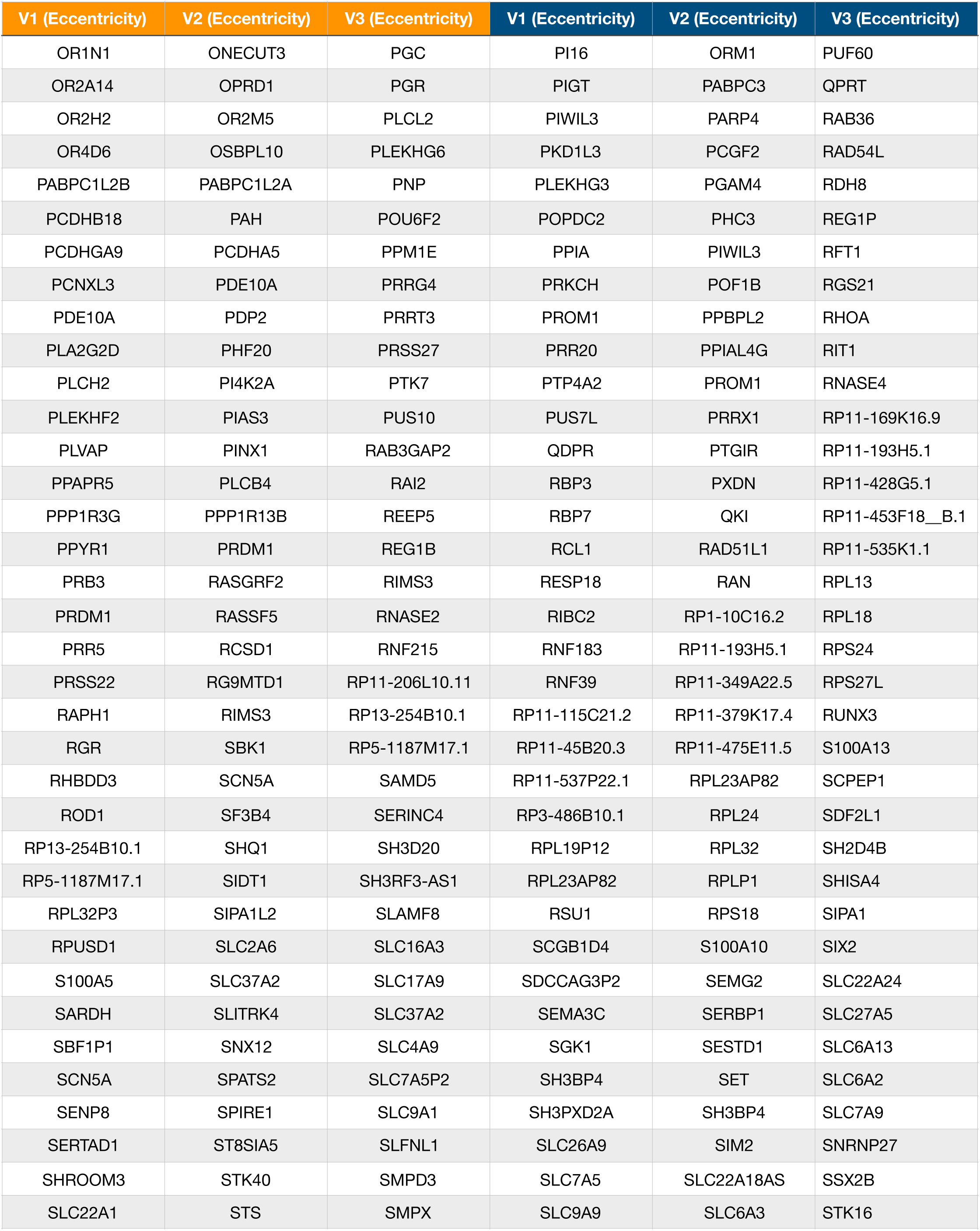
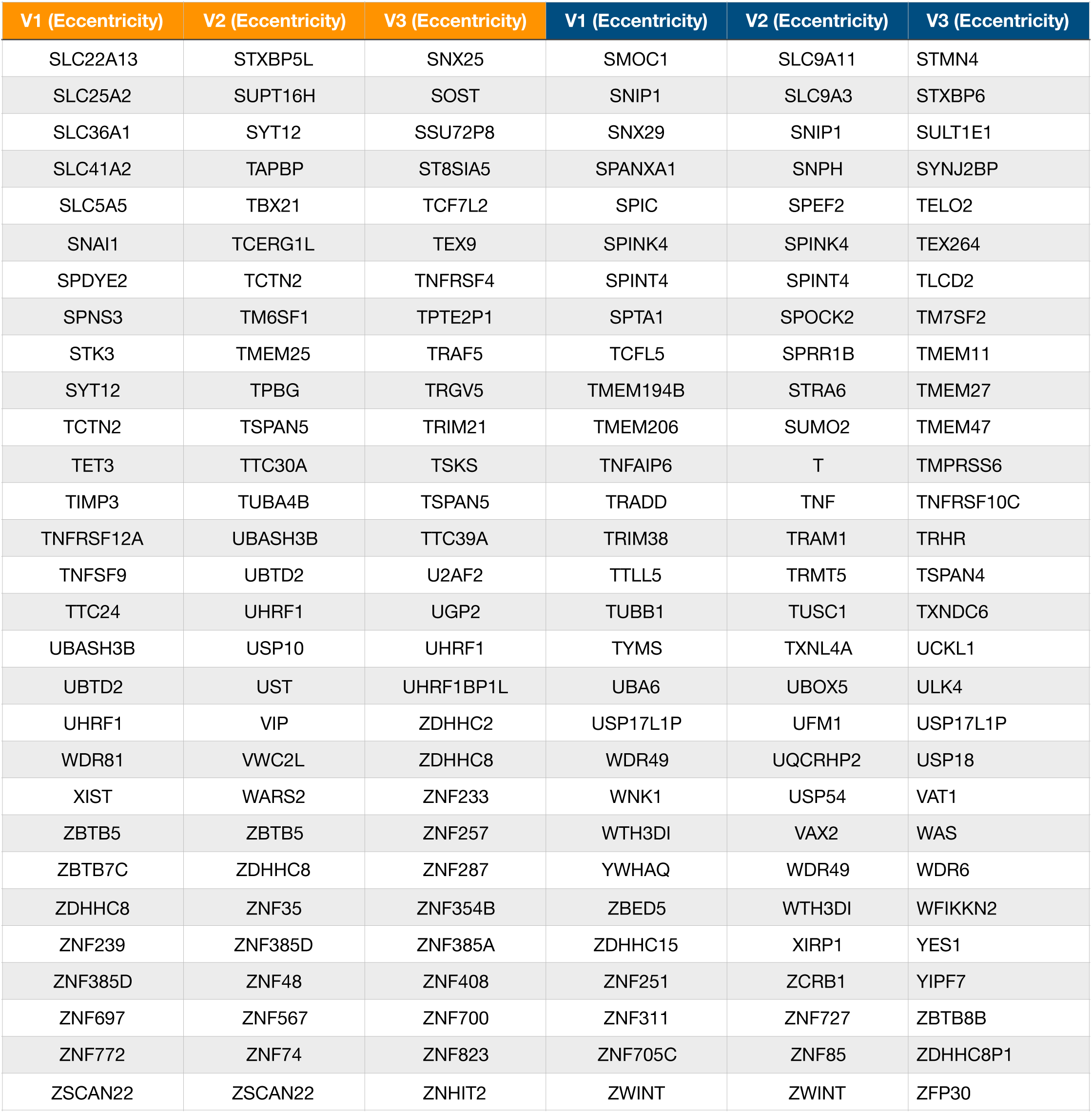
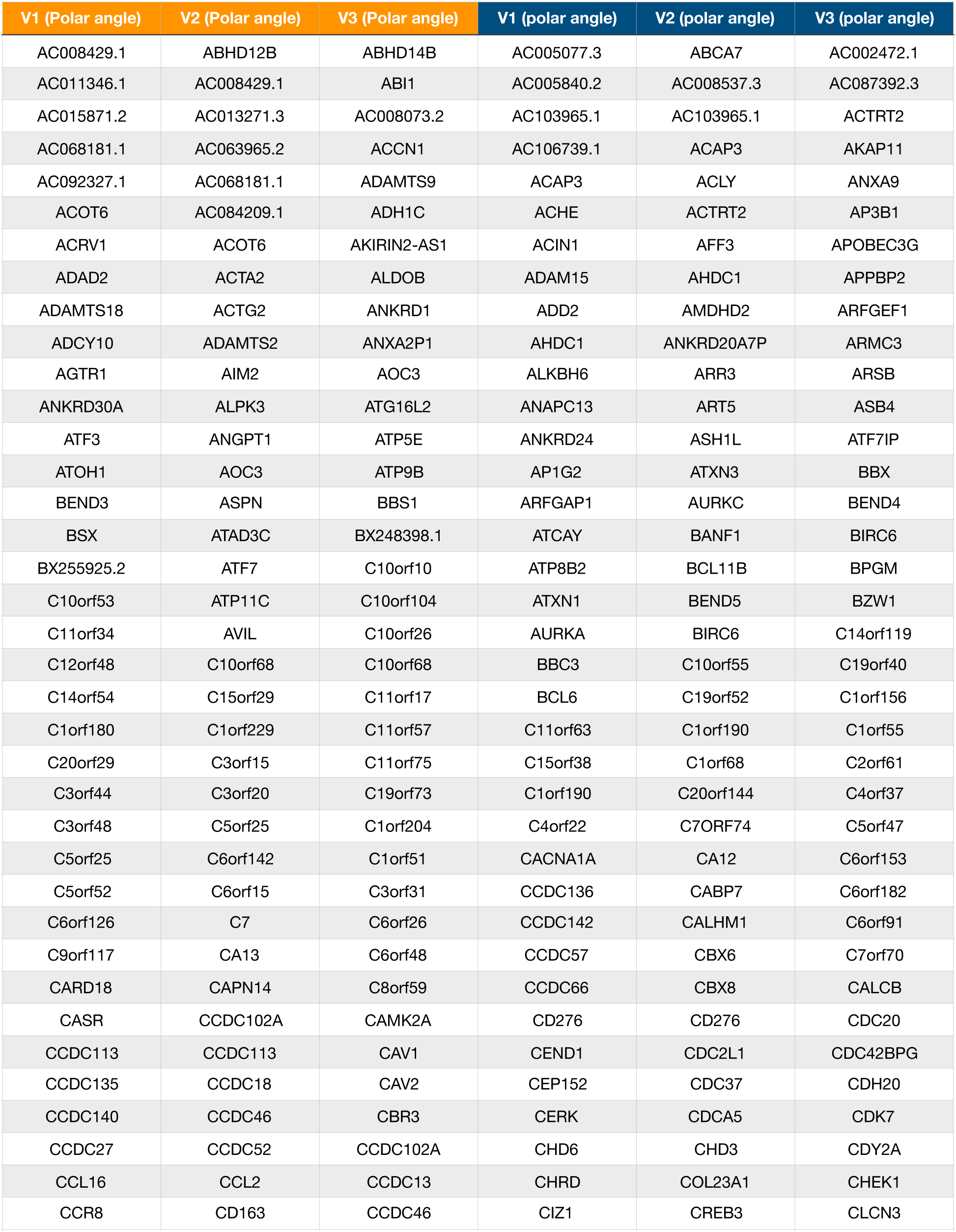
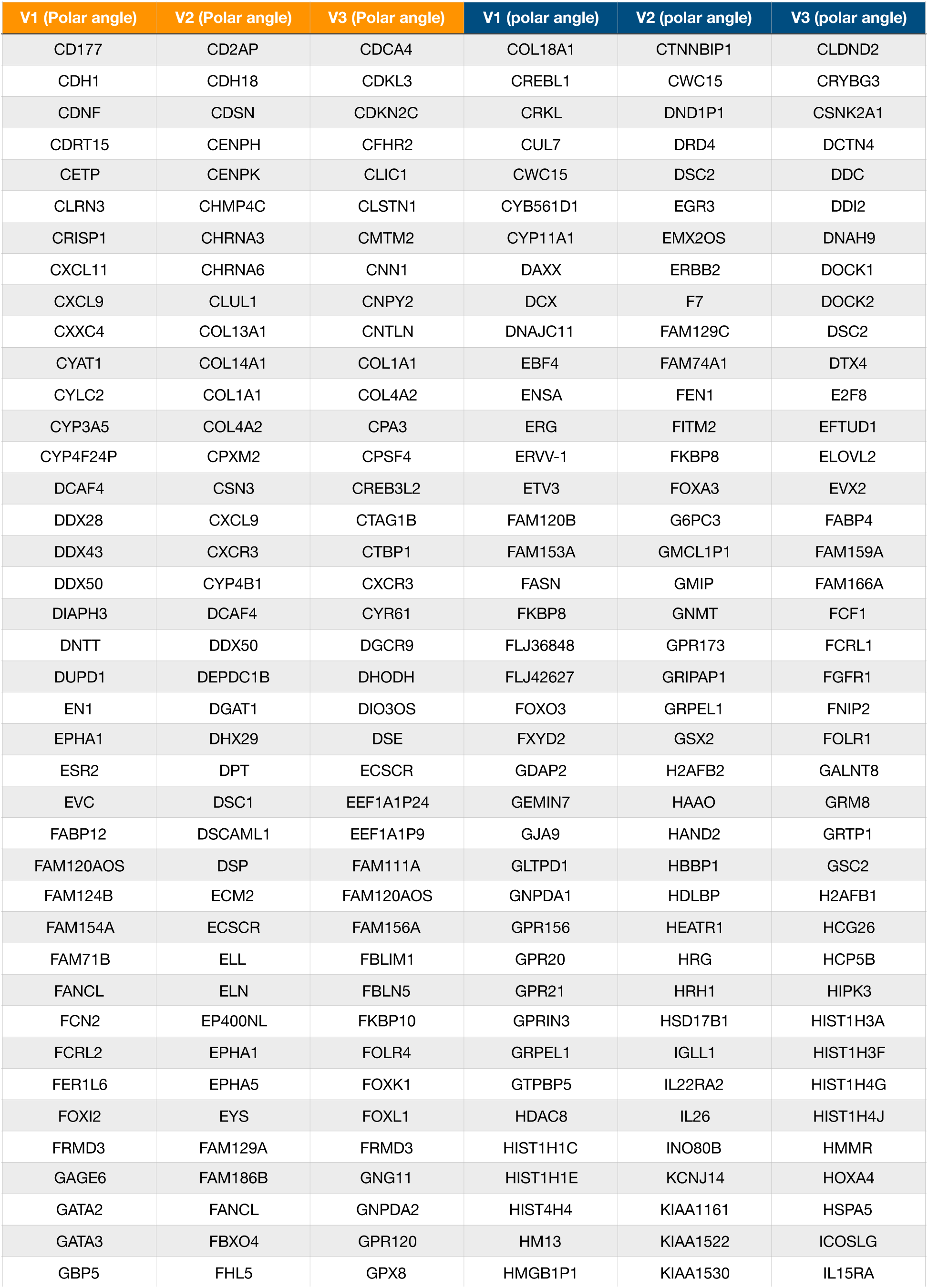
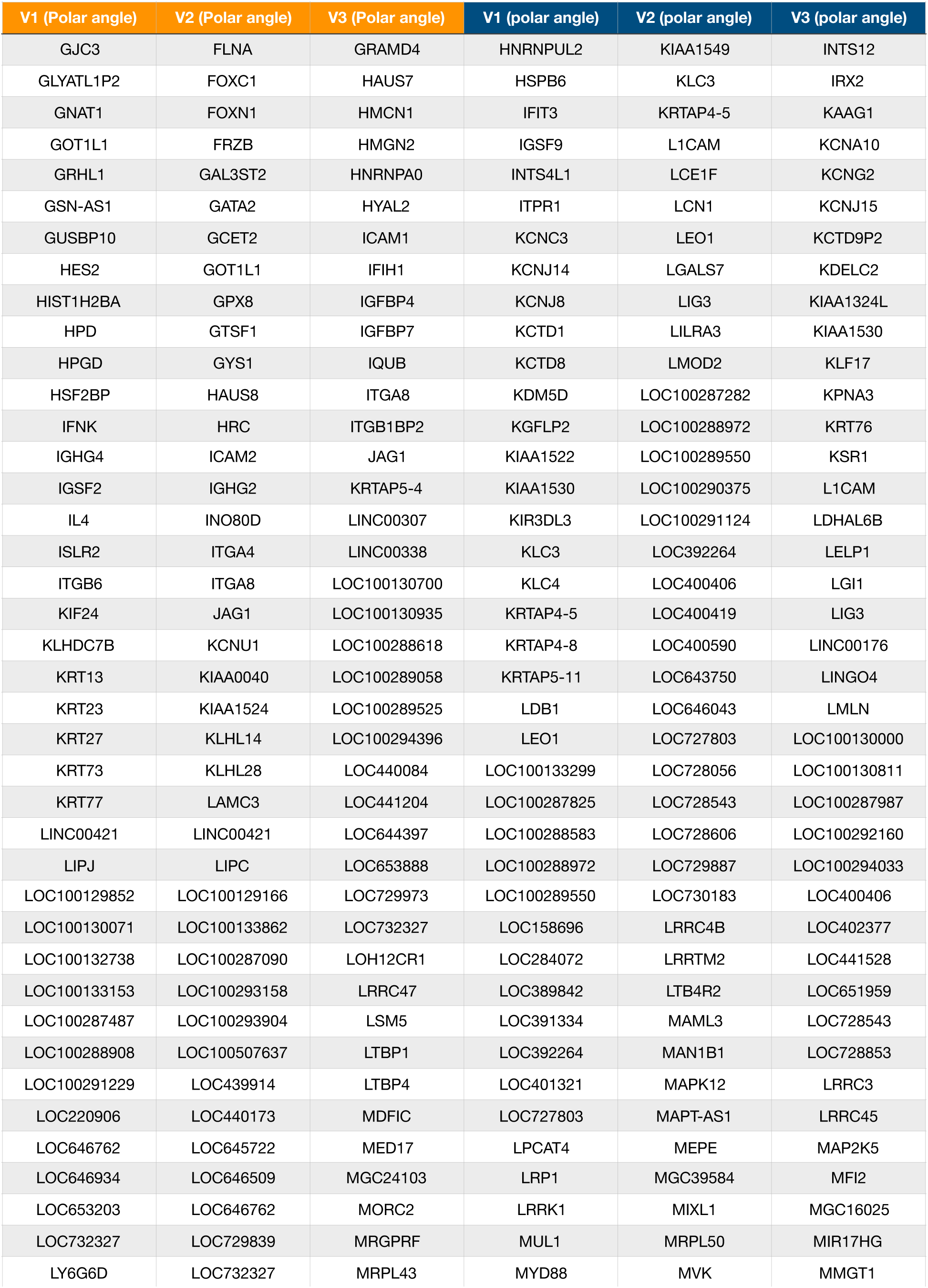
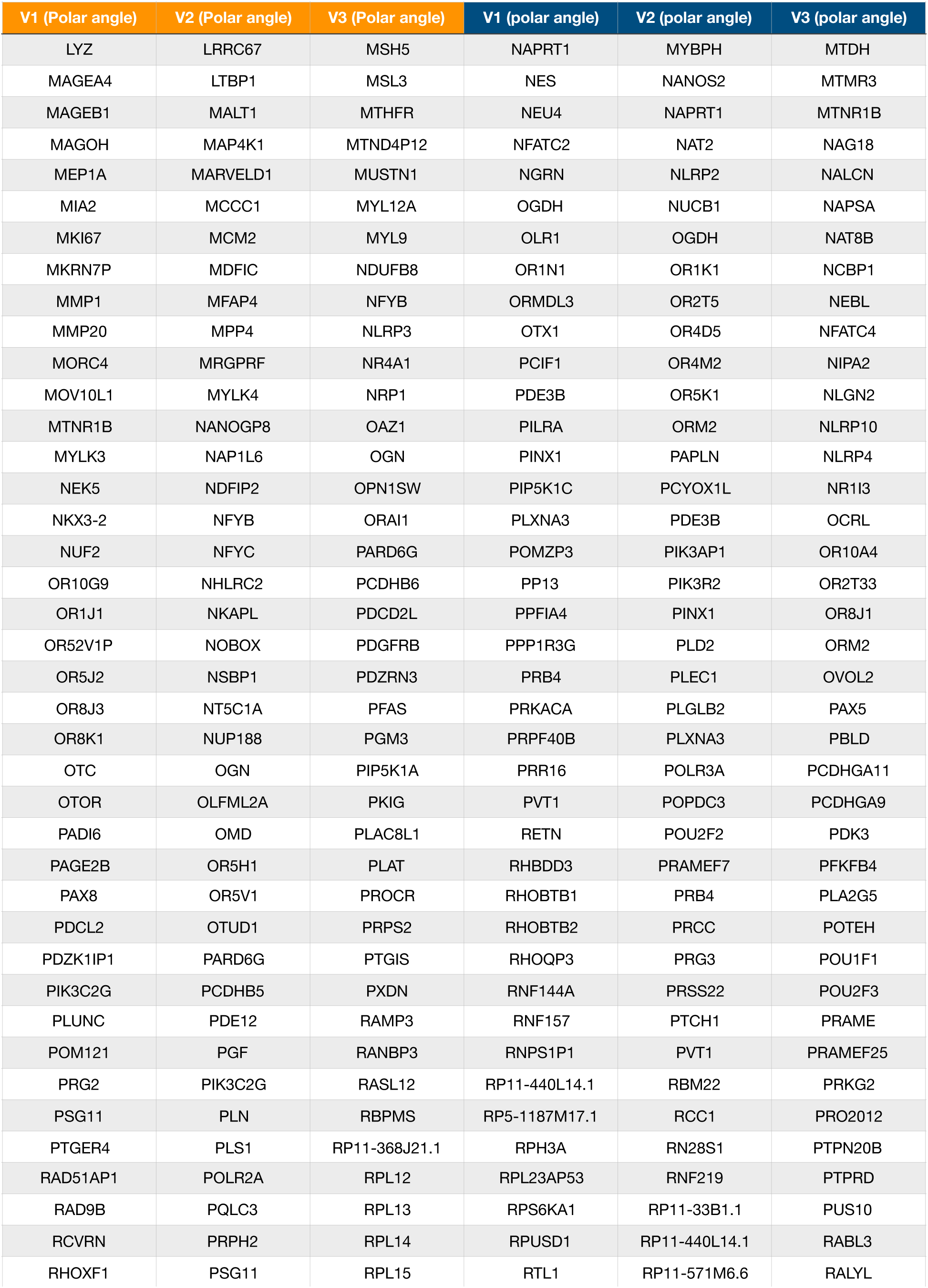
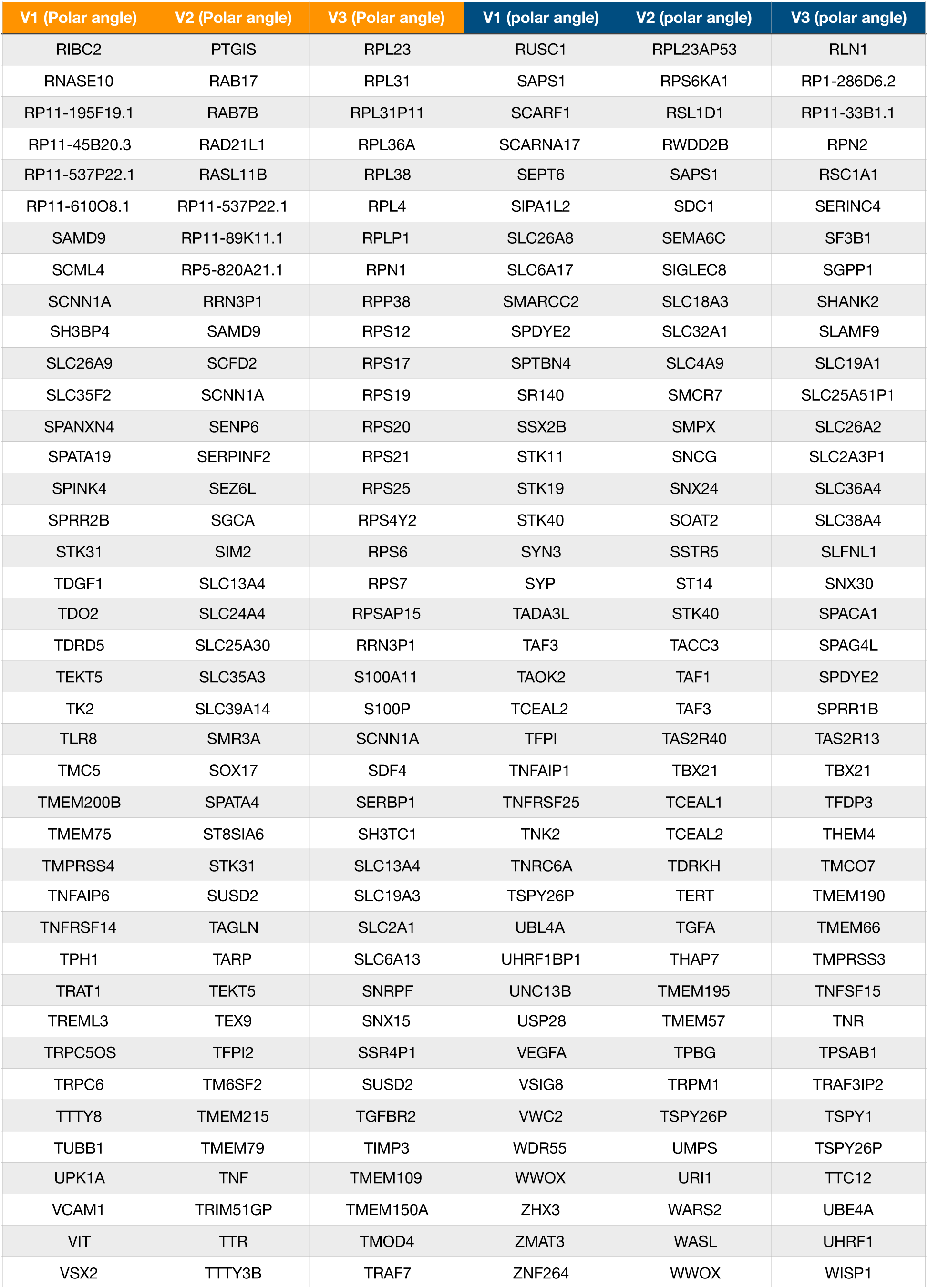
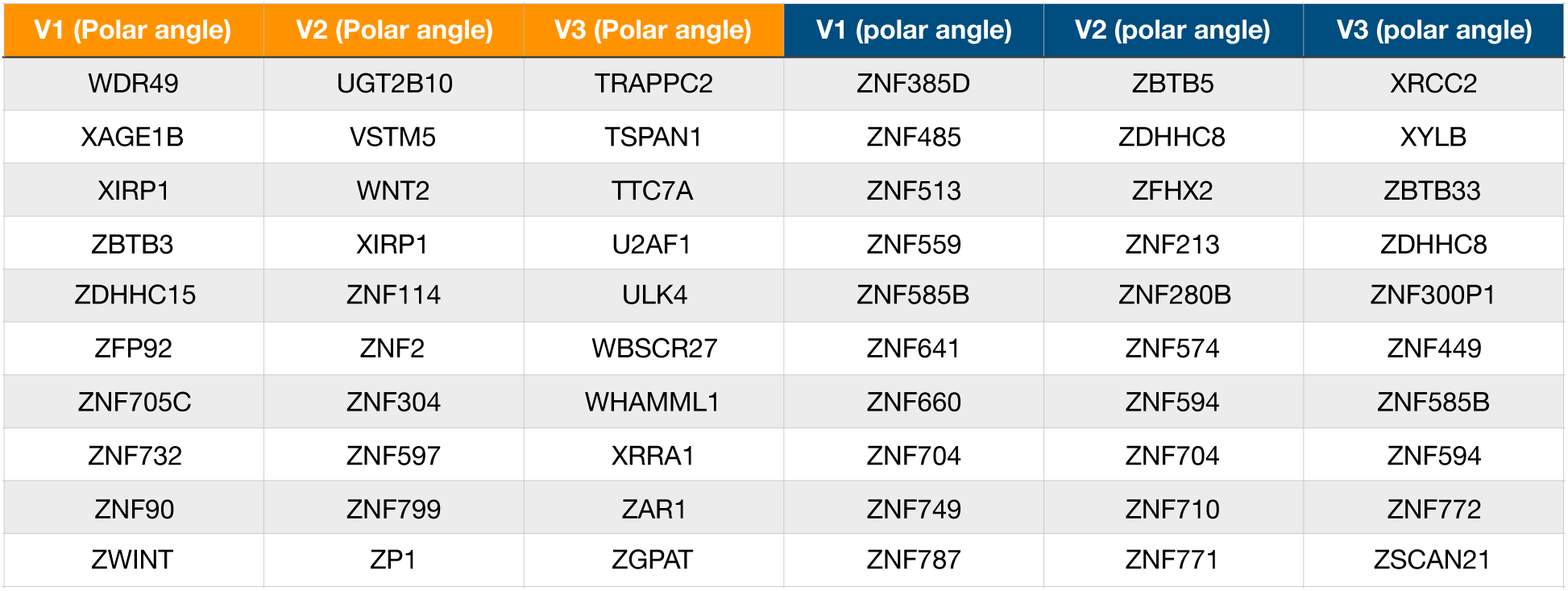
Genes correlated with eccentricity. Positively correlated genes are shown in orange, negatively correlated in blue. The top 1% of genes (ranked by -log of p-value) is shown for each field map.

**Supplementary Table 2.**

| **X chromosome**<br>**genes correlated**<br>**with eccentricity** |
| --- |
| **CXorf58** |
| **DCAF8L2** |
| **EDA** |
| **FLJ44635** |
| **GYG2** |
| **H2AFB2** |
| **HDAC8** |
| **HS6ST2** |
| **IQSEC2** |
| **OR13H1** |
| **PABPC1L2A** |
| **PGAM4** |
| **POF1B** |
| **PRRG3** |
| **RAI2** |
| **SLITRK4** |
| **SMPX** |
| **SNX12** |
| **SSX2B** |
| **STS** |
| **TMEM47** |
| **WAS** |

